# Revealing Shared Proteins and Pathways in Cardiovascular and Cognitive Diseases Using Protein Interaction Network Analysis

**DOI:** 10.1101/2023.08.03.551914

**Authors:** Melisa E. Zeylan, Simge Senyuz, Pol Picón-Pagès, Anna García-Elías, Marta Tajes, Francisco J. Muñoz, Baldo Oliva, Jordi Garcia-Ojalvo, Eduard Barbu, Raul Vicente, Stanley Nattel, Angel J. Ois-Santiago, Albert Puig-Pijoan, Ozlem Keskin, Attila Gursoy

**Affiliations:** Computational Sciences and Engineering, Graduate School of Science and Engineering, Koç University, Istanbul, Türkiye; Laboratory of Molecular Physiology, Department of Medicine and Life Sciences, Universitat Pompeu Fabra, Barcelona, Spain; Laboratory of Structural Bioinformatics (GRIB), Department of Medicine and Life Sciences, Universitat Pompeu Fabra, Barcelona, Spain; Laboratory of Dynamical Systems Biology, Department of Medicine and Life Sciences, Universitat Pompeu Fabra, Barcelona, Spain; Institute of Computer Science, University of Tartu, Estonia; Department of Medicine and Research Center, Montreal Heart Institute and Université de Montréal; Institute of Pharmacology, West German Heart and Vascular Center, University Duisburg-Essen, Germany; IHU LIRYC and Fondation Bordeaux Université, Bordeaux, France; Department of Neurology, Hospital Del Mar. Hospital Del Mar - Medical Research Institute and Universitat Pompeu Fabra, Barcelona, Spain; Department of Chemical and Biological Engineering, Koç University, Istanbul, Türkiye; Department of Computer Engineering, Koç University, Istanbul, Türkiye

**Keywords:** Network medicine, systems biology, crosstalk, cardiovascular diseases, cognitive diseases, vascular cognitive impairment, oxidative stress, protein protein interaction networks

## Abstract

One of the primary goals of systems medicine is detecting putative proteins and pathways involved in disease progression and pathological phenotypes. Vascular Cognitive Impairment (VCI) is a heterogeneous condition manifesting as cognitive impairment resulting from vascular factors. The precise mechanisms underlying this relationship remain unclear, which poses challenges for experimental research. Here, we applied computational approaches like systems biology to unveil and select relevant proteins and pathways related to VCI by studying the crosstalk between cardiovascular and cognitive diseases. In addition, we specifically included signals related to oxidative stress, a common etiologic factor tightly linked to aging, a major determinant of VCI. Our results show that pathways associated with oxidative stress are quite relevant, as most of the prioritized vascular-cognitive genes/proteins were enriched in these pathways. Our analysis provided a short list of proteins that could be contributing to VCI: DOLK, TSC1, ATP1A1, MAPK14, YWHAZ, CREB3, HSPB1, PRDX6, and LMNA. Moreover, our experimental results suggest a high implication of glycative stress, generating oxidative processes and post-translational protein modifications through advanced glycation end-products (AGEs). We propose that these products interact with their specific receptors (RAGE) and Notch signaling to contribute to the etiology of VCI.

## Introduction

Vascular Cognitive Impairment (VCI) is a heterogenous condition of vascular origin [1] and may lead to various forms of dementia, including Vascular Dementia (VD). VCI is generally a problem encountered in older adults, and the aging population has grown in the past century. Therefore, it is crucial to identify the biomarkers, therapeutic targets, and pathophysiological processes related to VCI. As VCI combines conditions characterized by both cardiovascular dysfunction and dementia manifestations, it is a complex and challenging syndrome to examine. The complexity of the pathologies also stems from their interconnected nature, which impacts brain function, vessel integrity, and heart activity. This interconnected nature makes the use of classical experimental approaches challenging.

We can improve our understanding of complex pathophysiological processes, such as diseases, by analyzing protein-protein interaction (PPI) networks [2]. PPI networks are constructed by integrating large-scale protein data into a network form, where nodes are the proteins and edges are the relations between them, such as physical and chemical interactions. The topology and measures of a PPI network are the main components that provide information about biological functions. Regarding their topology, biological networks, like PPI networks, are scale-free and contain “hub” nodes (nodes with high connectivity or degree), making them vulnerable to mutations at these hub nodes [3]. The degree centrality and betweenness centrality properties are two critical measures to analyze PPIs. While the degree centrality indicates how many connections a protein has in the network, betweenness centrality measures the importance of a protein regarding the number of shortest paths it lies on. Thus, betweenness centrality mediates the flow of information between nodes [4].

PPI network-based approaches are widely used to understand mechanisms underlying diseases [2, 5, 6]. However, PPI network-based approaches are still not extensively utilized to investigate the mechanisms related to VCI. Previous studies have demonstrated a significant association between vascular risk factors, cardiovascular diseases (CVD), and Cognitive Diseases (CD) [7, 8]. Considering this association, this study aims to identify key proteins and pathways involved in VCI by investigating the crosstalk between CVD and CD. Additionally, the role of Oxidative Stress (OS) in this crosstalk is investigated because OS increases during aging and is a condition known to be related to VCI [9], CVD [10], and CD [11]. OS increases due to disruption of the oxidant-antioxidant equilibrium [12] as a result of excessive production of Reactive Oxygen Species (ROS) by cells. ROS excessiveness can occur because of aging-related mitochondrial dysfunction [11] or amyloid β-peptide aggregates [13].

In this study, PPI networks related to CVD and CD phenotypes and a network related to OS were constructed. Here, we analyzed the crosstalk between CVD, CD, and OS-related PPI networks in two manners. First, we created a Global Network by merging (union) PPI networks and performed centrality analysis (Global Network Analysis). Second, we constructed the overlapping networks between CVD and CD subphenotypes, focusing on the not disease-associated proteins in the current literature (Overlap Network Analysis). These analyses suggest that OS plays a crucial role in the CVD-CD crosstalk and that DOLK, TSC1, ATP1A1, MAPK14, YWHAZ, CREB3, HSPB1, PRDX6, ATP13A2, LMNA are proteins highly dysregulated in this system. Also, we investigated the effect of OS in the different cell types in CVD and CD experimentally to determine the sensitivity to OS in heart, vessel, and brain cells. We discovered evidence that these cells have varying sensitivity to OS. Our experimental findings indicate that glycative stress plays a significant role in generating oxidative processes and post-translational protein changes via advanced glycation end-products (AGEs). Furthermore, our computational results support the existence of a crosstalk between AGE/RAGE and Notch Signaling pathways and propose that they are involved in VCI.

## Methods

### 1. Study Design

In order to assess VCI relevant proteins, the crosstalk between CVD and CD networks was studied. For this, a Disease-Phenotype Taxonomy was created. This contains several subphenotypes for CVD and CD. These subphenotypes were analyzed, and the ones with the highest seed protein count were selected to represent CVD and CD. Networks for these seven subphenotypes and OS were constructed with GUILDify. Additionally, a score is given by this tool for every protein in each network, signifying the relation of a protein with the subphenotype. These eight networks were first analyzed by Overlap Network Analysis. This analysis consists of constructing 17 overlap networks with the GUILDify tool. The constructed networks are CVD vs. CD, CVD vs. OS and CD vs. OS. Further, the highest scoring linker and non-seed proteins were searched for to assess novel relevant proteins. To find the scores in overlap networks, the average score of a protein with the given subphenotypes was calculated (Overlap Scoring Analysis). The top 3 highest scoring linker and non-seed proteins were closely analyzed to see if there are novel VCI biomarkers. The eight networks were next analyzed by merging them and constructing a Global network. Two scores were calculated for each protein on the Global network to assess the effect of OS (Global Scoring Analysis- **Fig. 1**): OS-excluded score and OS-included score. The Global Network was analyzed also topologically; however, as we also aim to assess the effect of OS, we categorized the proteins as whether they are OS related (present in the constructed OS network) or not. The OS proteins in the top 5 in the given centrality measures are determined as central OS proteins. We also refined these central proteins to the ones affected by at least 0.1. The central proteins were thus reduced from 22 to 12. We enriched these 12 proteins with (1) their CVD interactors, (2) their CD interactors, and (3) CVD&CD interactors. **Figure 2** demonstrates this methodology, and the rest of this section explains the methods in detail.

**Figure 1.**
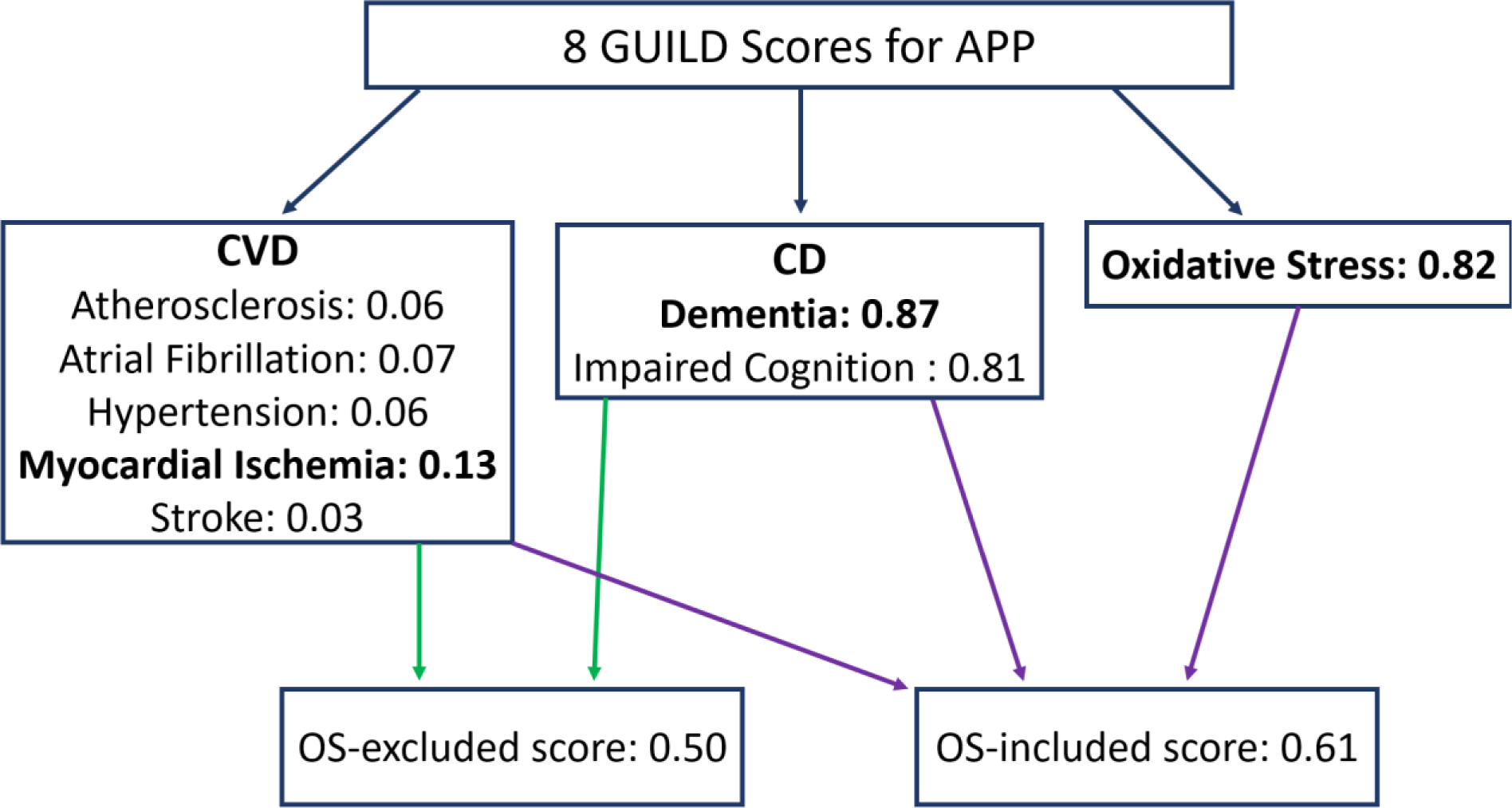
The flowchart for average GUILD score calculations. The average scores related to APP using Myocardial Ischemia, Dementia, and Oxidative Stress are given as an example. Figure demonstrates an example of how the prioritization is done for the APP protein. This calculation was done for all proteins on the GUILDify database (13090). As all the proteins on the GUILDify database have a GUILD score, we had 8 different GUILD Scores. The scores were categorized according to their Disease category, hence CVD, CD. Later we selected the phenotype with the maximum GUILD Score for its category. We obtained a possible crosstalk score for every protein by averaging the maximum CVD, and CD scores. We also included the GUILD Scores obtained from oxidative stress for comparison.

**Figure 2.**
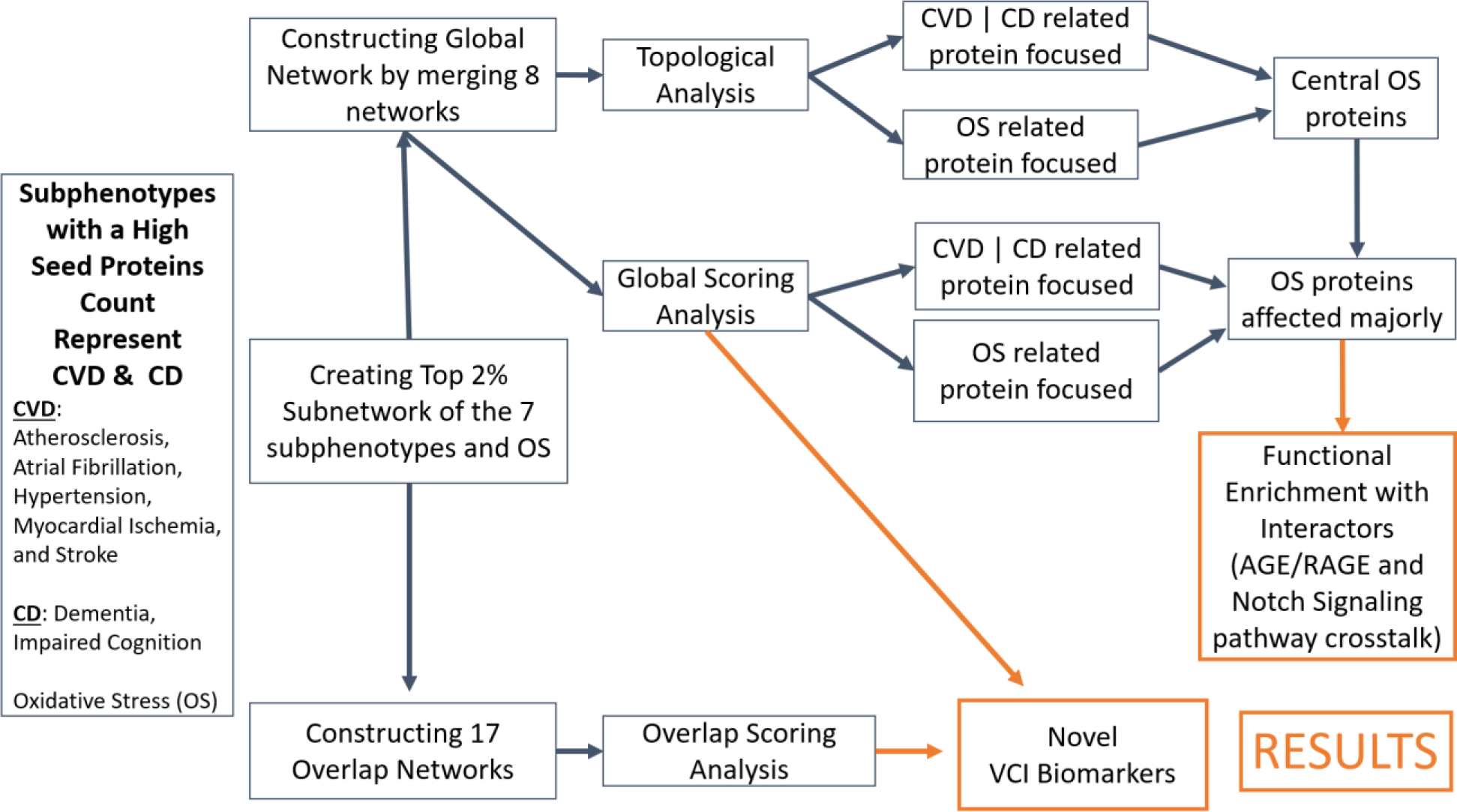
Research Methodology followed to understand the crosstalk between CVD and CD. The orange boxes represent results obtained.

### 2. Data Collection and Preparation

#### 2.1. Disease-phenotype taxonomy creation

Disease-phenotype taxonomy was created for CVD and CD. The taxonomy creation involved three steps. Initially, the first version of the taxonomy was derived from the Wikipedia category tree. Wikipedia category tree organizes Wikipedia articles into a hierarchical structure of categories and subcategories to assist users in navigating related content. Synonymous terms were added to each category using Princeton Wordnet [14]. Finally, the taxonomy was refined by two expert neurologists, Angel Ois and Albert Puig-Pijoan, who manually selected the most significant nodes and adjusted the structure. This <u>taxonomy</u> also included the synonyms of the diseases, if available. We chose three main branches for CVD and CD; and included all the sub-branches (subphenotypes) along with the synonyms of the selected disease-phenotypes.

#### 2.2. Selection of representative subphenotypes

The first step of detecting the crosstalk between CVD and CD was to select the subphenotypes from the disease-phenotype taxonomy that can be utilized to represent CVDs and CDs. As a short list of subphenotypes, we selected 80 subphenotypes, of which 30 are related to CD, and 50 to CVD.

Large counts of seed proteins highlight the abundance of information on literature about the relationship between proteins and the specific subphenotypes. Therefore, the selection of representative (of CVD, CD) subphenotypes was done according to the count of seed protein each subphenotype returned. Only 38 subphenotypes had more than one seed protein: seven subphenotypes belonging to CD and 31 to CVD categories. Furthermore, as we wanted to analyze the effect of OS in VCI, its seed protein count was also included (**Supplementary Fig. S1**).

We selected subphenotypes with the most significant count of seed proteins among different categories (CVD, CD) and discarded excessively general subphenotypes such as coronary artery disease. Subphenotypes representing CVD are Atherosclerosis, Atrial Fibrillation, Hypertension, Myocardial Ischemia, and Stroke. The subphenotypes representing CD are Dementia, and Impaired Cognition.

We selected the “impaired cognition” subphenotype instead of “Alzheimer” as the results must be specific for dementia and impairment as we want to analyze the earlier stages of cognitive degeneration, and also Alzheimer is considered as a unique particular case. Additionally, we do not want to bias our results with the stronger relation that Alzheimer’s has with aging. The “atrial fibrillation” subphenotype has the seventh largest seed protein count in the CVD instance, but it was chosen over “coronary artery disease” and “cardiovascular disease” because its network can reveal more precise mechanisms as it is a more specific term. Oxidative stress (OS) is relevant to both CVD and CD, so it was included for the crosstalk between CVD and CD.

### 3. Network Creation of the Representative Phenotypes

We applied the algorithms of GUILD using the GUILDify v2.0 Web Server [15]. GUILDify v2.0 is composed of two main components: BIANA [16] and GUILD. BIANA integrates the information of genes and proteins for protein-protein interactions and disease-phenotype associations, which are curated from literature. BIANA performs this part by extracting the information of genes and proteins from databases such as DisGeNet [17], UniProt [18] and OMIM [19], and for protein-protein interactions from MINT [20], DIP [21] and IntAct [22].

GUILD uses algorithms based on the guilt-by-association principle to prioritize/predict other protein/gene-disease associations. In GUILD, genes/proteins associated with a phenotype are named as “seed proteins” [16]. In the web server, a subnetwork with nodes achieving top scores (up to 1% or 2% of the network) and their interactions can be selected. These are called the Top 1% subnetwork and Top 2% subnetwork, respectively. Nodes that neither were seeds nor selected in the top but interact with two nodes of the selected network are named linkers. The web also allows the user to retrieve the top selected network including the linkers with highest score; and to calculate the functional enrichment of functions and pathways of the selected network. Finally, the intersection of two networks can also be retrieved and analyzed (for example to check the overlap between the results obtained for the subphenotypes of Stroke and Dementia). The algorithm of NetCombo of GUILD was applied to perform the prioritization. This is a combination of NetScore, NetZcore and NetShort algorithms, averaged after normalization. The tissue type was selected as “all”. A total of eight PPI networks were retrieved from GUILDify v2.0 by selecting the top 2%, including their linkers and seeds.

### 4. Construction and Analysis of the PPI Networks

#### 4.1. Constructing 17 overlap networks and overlap scoring analysis

In order to provide a different perspective on the proteins on the crosstalk of CVD and CD, we constructed 17 phenotype pair overlap networks with GUILDify. For each of the overlap networks (**Supplementary Table S1, Supplementary Table S2**), the GUILD scores of the linker and non-seed proteins were calculated. As GUILDify does not provide scores for the overlap, we calculated the scores by taking the average of the scores of subphenotype 1 and subphenotype 2. This score will be referred to as the “overlap score”.

#### 4.2. Global network construction, global scoring analysis and functional enrichment

We created a Global Network by merging the networks of the selected 7 subphenotypes and OS. The Global Network was visualized with Cytoscape. The node label sizes are correlated to their degree such that for degree 0 to 19 the labels are not shown. For degrees from 21 to 41 the label size is 25, from 42 to 90 it increases to 30, and from 90 to max it increases to 35. The Global Network was analyzed topologically by degree and betweenness centrality. We categorized the top scoring proteins whether they are OS related or not. The network analysis is performed with the python module NetworkX version 2.8.8 [23]. We detected 22 central OS proteins. However, to refine these 22 proteins to the most affected ones by OS, we performed the Global Scoring Analysis.

In the Global Scoring analysis for each protein, we selected the highest GUILD score of a category and averaged the GUILD Scores of different categories. We performed this in two settings: first one is to average CVD and CD related GUILD Score for each protein, and the second is to average CVD, CD, and OS. **Figure 1** demonstrates this flow of creating two different scores for each protein. Here, we aimed to analyze the effects of OS related proteins on the crosstalk by creating two average GUILD scores to prioritize proteins on the crosstalk. An example of the prioritization is given in **Fig. 1**.

The 22 proteins were refined to 12 by selected the proteins that are affected 0.1 when the OS GUILD score is included in the average. These are the central OS proteins. Lastly, in order to detect the crosstalk pathways, these 12 proteins are enriched in 3 different settings. First with their CVD interactors present in the Global Network, secondly with their CD interactors present in the Global Network and lastly with their CVD&CD interactors (both). For all the functional enrichment analysis g: Profiler was used.

### 5. Experimental Procedures

#### 5.1. Cell lines

All the cell lines were obtained commercially and grown according to their special requirements plus 5-15% fetal bovine serum (FBS) and 1% streptomycin/penicillin at 37°C in a humidified atmosphere containing 5% CO2.

#### 5.2. Murine cortical primary cultures

Cortex were isolated from 18-day-old CB1 mouse embryos following the procedure approved by the Ethics Committee of the Institut Municipal d’Investigacions Mediques- Universitat Pompeu Fabra. Brain samples were dissected, trypsinized and cells were isolated and seeded in phenol-red-free Dulbecco’s modified Eagle’s medium (DMEM; Sigma) plus 10% horse serum into 1% poly-D-Lysine coated coverslips (5x10^4^ cells/cover). After 120 min, the medium was removed and neurobasal medium containing 1% B27 supplement (Gibco BRL) plus 100 units/mL penicillin and 100 mg/mL streptomycin. 2 µM cytosine arabinoside (Sigma) were added at day 3 for 24 h to avoid glial proliferation. Cultured cortical neurons were used for the experiments on day 7.

#### 5.3. Cell viability assays

Cells were seeded in 96-well plates at a density of 2,5x10^4^ cells/well. Cells were treated win increasing concentrations of H_2_O_2_ or MG and incubated 22 h. 10% of 3-(4,5-dimethylthiazol- 2-yl)-2,5-diphenyltetrazolium bromide (MTT) stock solution (5 mg/mL) was added per well and the reaction was stopped with 120 μL of DMSO after 2 h. MTT reduction was determined in a plate reader spectrophotometer at 540 and 650 nm. Control cells were assumed as 100%.

#### 5.4. Immunofluorescence study of MG treatment

Cells were seeded on poly-L-lysine coated coverslips in 24-well plates at a density of 5x10^4^ cells /well. After 12 h, the medium was removed and Ham’s F12 without FBS was added to the wells. Then cells were treated with MG. After 24h, cells were fixed with 4% paraformaldehyde and permeabilized with 0.1% Triton X-100. Cortical primary cultures were immunostained with 1:100 rabbit Caspase-3 Antibody (Ab; Cell Signaling, Beverly, USA), 1:1000 rabbit Tuj-1 Ab (Covance, San Diego, CA, USA), and Topro (Life Technologies, Carlsbad, CA, USA); and 1:2,000 Alexa 555- or 488-bound as secondaries Ab (Sigma, St. Louis, USA) at room temperature. Coverslips were mounted and analyzed using a Leica TCS SP confocal microscope and analyzed with Leica confocal software.

## Results

We used PPI networks to assess the crosstalk between CVD and CD and unveil the biological functions associated with VCI. Eight PPI networks related to CVD, CD subphenotypes and OS were constructed. Each protein in each network had a score indicating its association with a subphenotype. These scores were used for two different scoring analyses, and these were used to find novel proteins and central proteins. Also, each protein in each network has seed, linker, or non-seed properties (see Methods Section 2-3 for details). These eight PPI networks were analyzed by (1) Overlapping CVD&CD, CVD&OS, and CD&OS networks and (2) Merged Global Network. (See Methods Section 4 for details). From both analyses we obtained a number of potentially novel VCI-related proteins. The topological and scoring analyses of the Global Network indicated a crosstalk between AGE/RAGE and Notch signaling pathways, which could be related to VCI pathology. And lastly, the sensitivity of heart, vessel, and brain cells to OS was experimentally determined (See Methods Section 5 for details).

### Analyses of overlapping networks predicted novel VCI-relevant proteins

Ten overlapping networks were constructed by overlapping the selected/prioritized CVD and CD subphenotypes (**Supplementary Table S1)**. An example is shown in **Fig. 3** for the overlap of genes/proteins of Atherosclerosis and Dementia subphenotypes. Each protein in the network was identified either as a linker, non-seed, or seed protein. Linker and non-seed proteins are potentially novel proteins that can be related to VCI. This is because while seed proteins are curated from literature, the others come from the prioritization of the GUILDify algorithm. Additionally, each protein in the overlapping network was scored. The linker and non-seed proteins with the highest top 3 overlap scores were included in **Supplementary Table S1**. Interestingly, most of these proteins were linker proteins. Out of the 30 proteins given in **Supplementary Table S1**, 22 were unique, and 5 of them (VEGFA, ATXN1, DOLK, FLNA, GJB2) were present in more than one overlapping network. Interestingly, 14 of them were also associated with OS (i.e., they are associated or predicted to be associated with OS, they are obtained from the OS network); these were: HMGB1, TSC1, REL, RELA, SH2B1, ATP1A1, COL5A1, FLNA, SDHB, LMNA, GJB2, DOLK. Similarly, **Supplementary Table S2** shows the analyses of the overlaps between the seven subphenotypes of CVD (5) and CD (2) with OS. 13 out of 21 proteins with the highest top 3 overlap scores were linker proteins of the original subphenotype networks. Furthermore, despite the lack of shared proteins between these overlapping networks, two proteins, REL and ATP13A2, are also present in the table of top genes/proteins with overlapping scores between CVD and CD.

**Figure 3.**
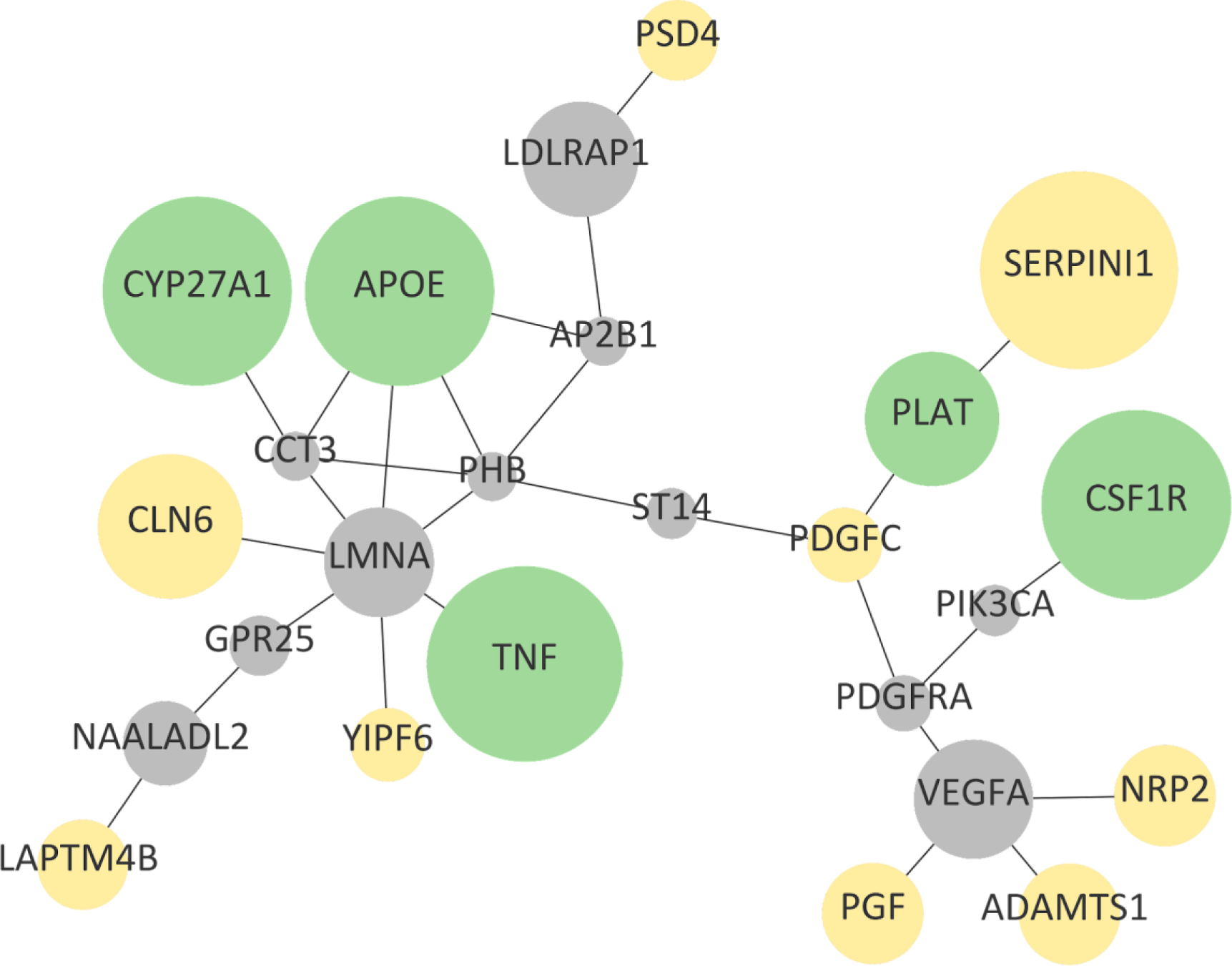
Overlap network between the prioritized proteins associated with subphenotypes Atherosclerosis and Dementia. The size of the nodes (proteins) is correlated with the overlap score (i.e. the size of the highest overlap score is four times the lowest). Edges show protein-protein interactions between seed proteins (in green), linkers (in gray) and non-seeds with top scores of associations with their respective subphenotypes (in yellow).

### Global Network Analysis focused on crosstalk pathways

**Figure 4** shows the Global Network formed by the sum of all subphenotype networks and OS networks. The Global Network was analyzed by topological measures of network-centrality (degree centrality and betweenness centrality, see Methods section). The top 5 proteins with the highest degree of centrality and/or betweenness centrality were defined as central proteins. To measure the effect of OS in the Global Network, we performed two analyses: 1) Assigning and comparing OS-included and OS-excluded scores to every node, and 2) Topological analysis to find central OS proteins and central non-OS proteins.

**Figure 4.**
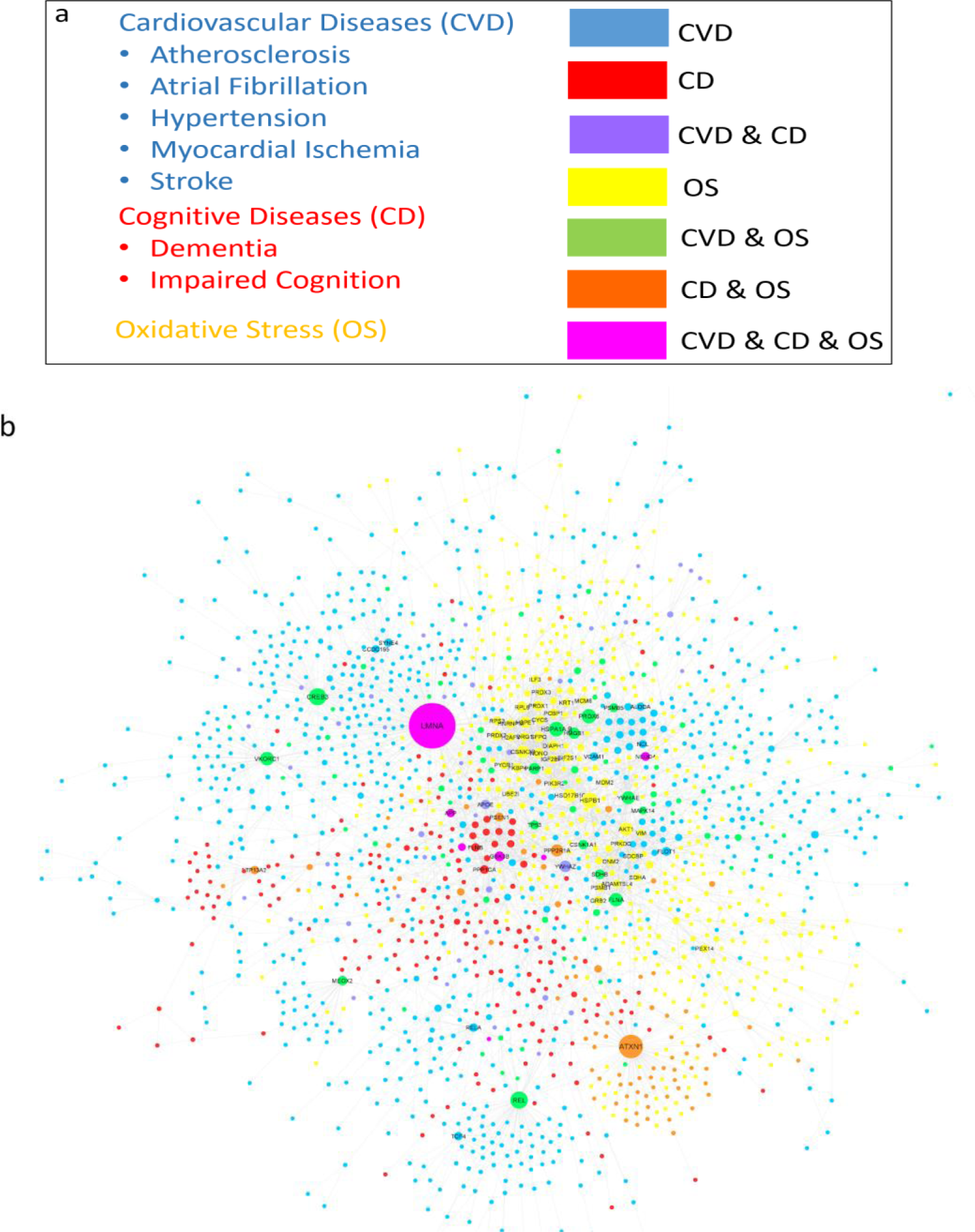
**(a)** Labels of the global network nodes. The nodes are colored according to the disease originating from their subphenotype network or shared by more than one. **(b)** Global network. Node sizes are correlated with their degree (see Methods for details). A prefuse force-directed layout with edge-betweenness was utilized to draw the network with Cytoscape [24] and the overlapped nodes were removed. To facilitate interpretation the node label sizes are correlated with their degree, also represented by their gene symbol.

Two scores were calculated for each node in the Global Network: a CVD-CD average score (OS-excluded) and a CVD-CD-OS average score (OS-included score). **Supplementary Figure S2** demonstrates the distribution of the calculated scores of the proteins in the Global Network. The difference between the OS-excluded and OS-included scores was always smaller than 0.2, with variations around 0.1 (see **Supplementary Fig. S2**). Around 33% of proteins of the Global Network showed a difference between OS-excluded and OS-included scores: 255 proteins out of 1796 showed an increase of about +0.1 and 238 a decrease of about -0.1, while the rest were not affected (or the difference was not noticeable). Some of these proteins were further investigated for VCI relation, annotated by genes: HMGB1, SOD2, MAPK14, and JAK2.

The Global Network was topologically analyzed (**Supplementary Table S3**). In the topological analysis of the Global Network, the average shortest path length was found to be 4.97. Furthermore, the clustering coefficient was found to be 0.69, which can be considered a high value, as the maximum value is 1. According to the centrality measures LMNA, ATXN1, REL, FLNA, CREB3, HSPB1, and YWHAZ were the central proteins. Interestingly, except for YWHAZ, all other central proteins were from the OS network. To determine the role of OS, central proteins of the Global Network were further analyzed based on whether they were associated with OS (i.e., obtained from the OS network) or not, distinguishing between central OS non-proteins and central OS proteins, respectively.

The central non-OS proteins found were: ALDOA, RELA, SMAD1, FLOT1, DLG4, SYNE4, YWHAZ, FLNB, SMAD4, APOE, VCAM1, PPP1CA, CAV1, FN1, EWSR1, TCF4. The proteins produced by these genes had a total of 215 interactors in the Global Network. Of them, 174 interactors were only associated with CVD, 31 only with CD, and ten were common to CVD and CD subphenotypes. To better understand the biological processes involved, functional enrichment of each set of these central proteins and their CVD and CD interactors (from the Global Network) was found. **Supplementary Table S4** shows the functional enrichment of the non-OS central proteins and their interactors.

To detect the pathways and processes related to the effect of OS in the crosstalk, we determined the central OS proteins (**Table 1**). **Table 1** has a total of 22 unique proteins, 12 out of them with a difference of +/- 0.1 between OS-excluded and OS-included scores (produced by genes HSPB1, AKT1, PRDX6, FLNA, VKORC1, ATXN1, PSEN1, PRKN, ATP13A2, NEDD4, APP, and LMNA). **Supplementary Table S5** lists the 390 interactors of these 12 proteins (250 from CVD subphenotype networks, 160 from CD, and 20 of them common to both CVD and CD), and **Fig. 5** shows the enriched pathways (AGE/RAGE and Notch signaling) for CVD and CD interactors, respectively. Interestingly, our findings suggest that oxidative-glycative processes are particularly relevant since advanced glycation end products (AGE), which are quite relevant in diabetes, also play a key role in VCI by their interaction with Notch signaling, a regulator of growth and maintenance of different tissues that are particularly important in blood-brain barrier integrity.

**Figure 5.**
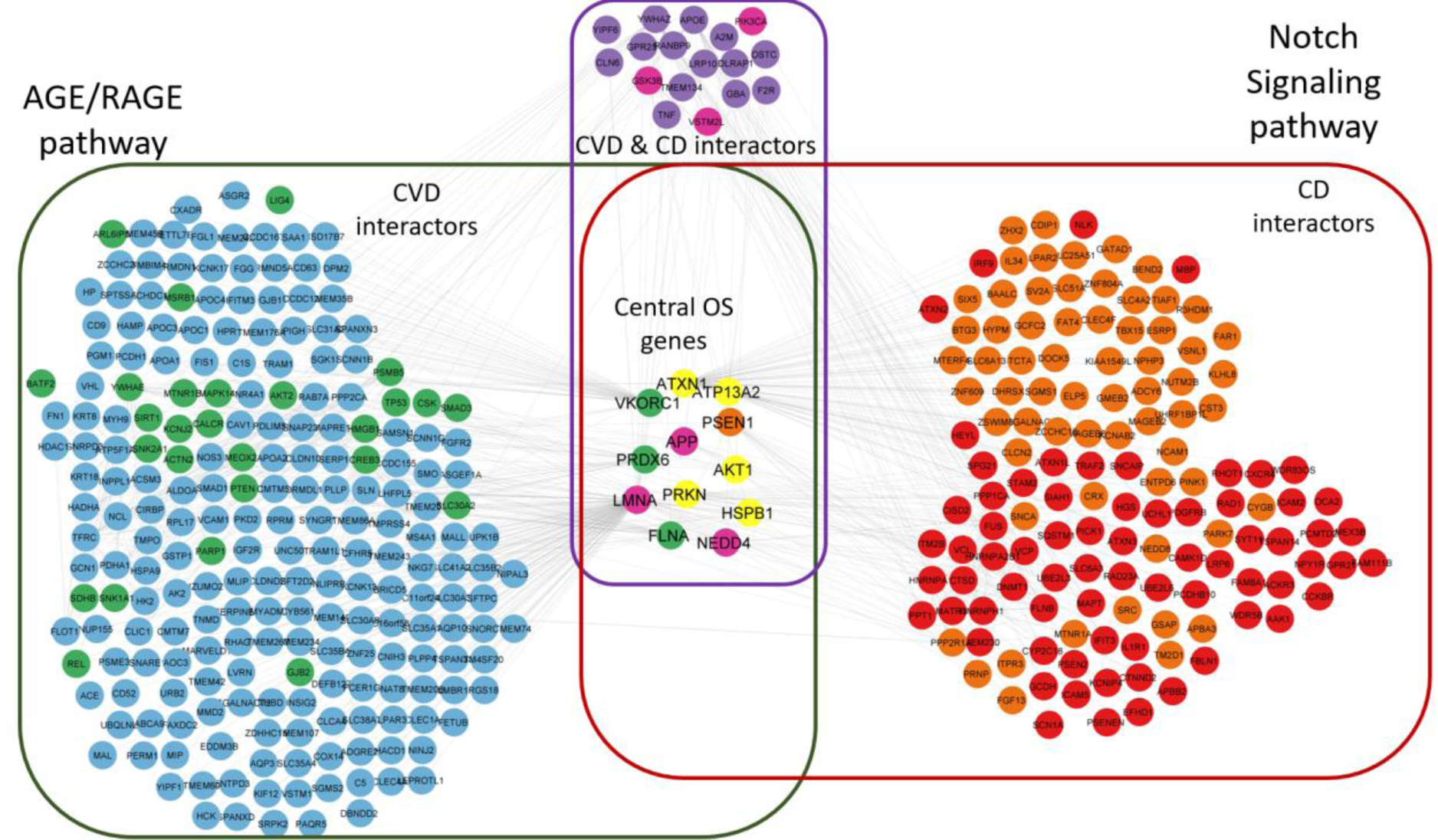
Pathway enrichment of the subnetwork formed by the central OS proteins and their interactors: The AGE/RAGE for CVD (green box), the Notch signaling for CD (red box).

**Table 1.** Central proteins of the Global Network^1^.

| Protein Category | Degree Centrality | Betweenness Centrality |
| --- | --- | --- |
| OS | • **HSPB1**<br>• HSD17B10<br>• **AKT1**<br>• CSNK2B<br>• HSPE1 | • **AKT1**<br>• **HSPB1**<br>• HSD17B10<br>• UBE2I<br>• CSNK2B |
| CVD & OS | • REL<br>• CREB3<br>• **PRDX6**<br>• **FLNA**<br>• YWHAE | • REL<br>• **FLNA**<br>• CREB3<br>• **VKORC1**<br>• YWHAE |
| CD & OS | • **ATXN1**<br>• PPP2R1A<br>• **PSEN1**<br>• **ATP13A2**<br>• **PRKN** | • **ATXN1**<br>• PPP2R1A<br>• **PRKN**<br>• **PSEN1**<br>• **ATP13A2** |
| CVD & CD & OS | • **LMNA**<br>• GSK3B<br>• **NEDD4**<br>• **APP**<br>• COPS5 | • **LMNA**<br>• GSK3B<br>• **APP**<br>• **NEDD4**<br>• COPS5 |

<sup>1</sup> Selected central proteins with the top 5 highest measures of degree centrality and betweenness centrality from CD and CVD subphenotype networks and/or the OS network are indicated by their gene names. Those with a differential score between OS-excluded and OS-included scores around +/- 0.1 are highlighted in bold.

### The effect of oxidative and glycative stress varied on heart, vessel, and brain cells

We studied the effect of OS in the different cell types involved in CVD and CD experimentally (**Fig. 6**) to assess the sensitivity to OS in heart (cardiofibroblasts), vessels (endothelial cells and vascular myocytes), and brain cells (neurons and microglia). We found evidence that, under the same conditions, these cells have differential sensitivities to OS, with vascular cells being the most sensitive (**Fig. 6B**). Endothelial cells showed statistically significant reduced viability at 10 µM and 50 µM H_2_O_2_ (p<0.001 for both concentrations) and human aortic vascular smooth muscle cells at 50 µM H_2_O_2_ (p<0.001). However, human neuroblastoma cells (**Fig. 6C**) only showed decreased viability at 100 µM and 500 µM H_2_O_2_ (p<0.01 and p<0.001 respectively). Murine microglial cells did not show any statistically significant effect in their viability even when they were challenged with 500 µM H_2_O_2_. In addition, cardiofibroblasts (**Fig. 6A**), cells believed to play a major role in atrial fibrillation [25, 26], were quite resistant to OS since just 500 µM H_2_O_2_ was cytotoxic for these cells (p<0.001).

**Figure 6.**
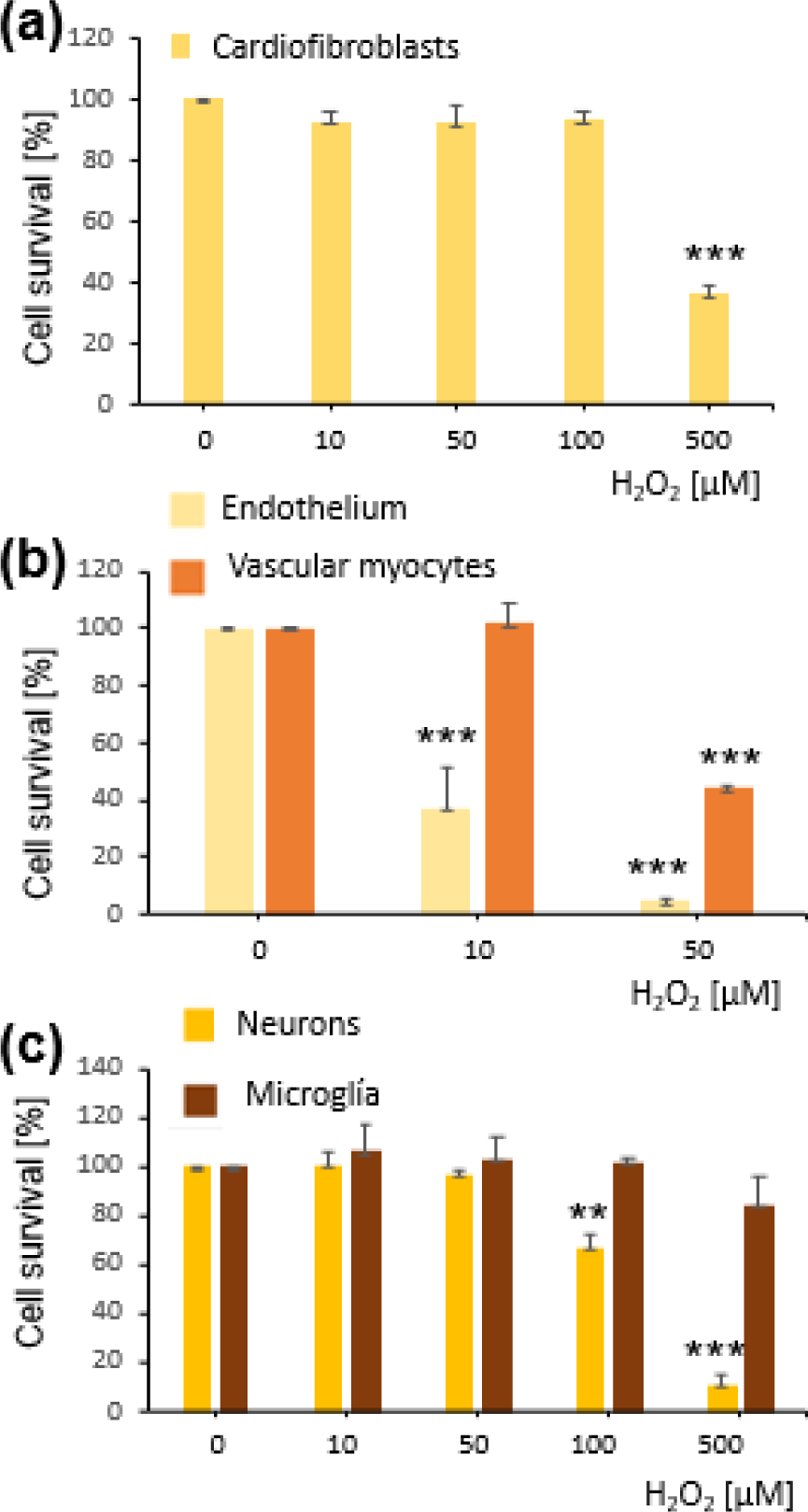
OS effect on hearth (**a**), vascular (**b**) and brain cells (**c**). Cell lines of human cardiofibroblasts, HCF, human umbilical vein endothelium (HUVEC), human aortic vascular smooth muscle cells (HA-VSMC), human neuroblastoma (SH-SY5Y) and murine microglia (BV2) were challenged with increasing concentrations of H_2_O_2_ for 24 h. Cells viability was assayed by MTT reduction. Data are the mean ± SEM of 3-7 independent experiments performed by triplicate. ** p<0.01, *** p<0.001 *vs* the respective controls by ANOVA plus Tukey-Kramer Multiple Comparisons Test.

Furthermore, the relevance of the oxidative-glycative stress was assayed in vitro using methylglioxal (MG) (**Supplementary Fig. S3**), which is able to induce protein glycation by post-translational modifications of specific amino acids [27, 28]. We found that 250 µM MG shows statistically significant toxicity for mature murine cortical neurons, as measured by cell viability assays and caspase 3 activation.

## Discussion

Our study aims to uncover key proteins and pathways related to VCI by exploring the crosstalk between CVD and CD and addressing the involvement of OS in this crosstalk. While network analysis showed OS has a substantial effect in the crosstalk, cell viability experiments demonstrated that the effect of OS varies in different cell types. Furthermore, we computationally hypothesized that the crosstalk between AGE/RAGE and Notch signaling pathways could be crucial for VCI. Experimental results revealed glycative stress damages neurons.

### Evaluation of the Identified Proteins’ Relation to VCI

In relation to identifying novel VCI-relevant proteins, several proteins had the potential to be related to VCI. While some of these proteins were identified using overlap or Global Network analysis, others were identified through both. 16 proteins from **Supplementary Table S1,** which had at least 0.4 overlap score and were related to OS, were investigated for VCI relation. Eight of these proteins also excelled in the Global Network analysis. Furthermore, 23 proteins were selected to be potential VCI-relevant proteins from the Global Network analysis. Eight of these were also present in the top results of the overlap network analysis. **Supplementary Table S2** aimed to demonstrate crosstalk proteins by investigating the linker and non-seed proteins with the pairwise analysis between CVD and OS vs. CD and OS. It is important to note that 17 of these proteins have at least a 0.43 overlap score, being relatively high as this score is based on an average value.

As the aim was to identify novel relevant proteins involved in VCI, we searched the literature by their relationship with “vascular cognitive impairment”, “vascular dementia” and “cerebrovascular disease” to determine which of these proteins are novel VCI-relevant proteins. Furthermore, if a protein was not previously recognized as VCI-related, a relation with CVD and CD was searched simultaneously or individually. **Supplementary Table S6** summarizes the proteins and their association with CVD, CD, or VCI. Proteins with no known association with VCI in the literature but known associations with CVD and CD in different studies were examined as possible novel VCI-relevant proteins. From **Supplementary Table S6**, it was deduced that among the proteins found crucial only in Overlap analysis; VEGFA, GJB2, COL5A1, SDHB, and SLC30A10 had already been reported as VCI-related proteins [29–33]. DOLK, TSC1, and ATP1A1 were related to both CVD and CD [34–39]; however, no research demonstrated it as VCI related.

Of the proteins found crucial only based on Global Network analysis, SOD2, JAK2, APOE, ALDOA, VCAM1, AKT1, APP, PSEN1, NEDD4, and PRKN were previously already related to VCI [40–50]. Relations to both CD and CVD were found for MAPK14, YWHAZ, CREB3, HSPB1, and PRDX6 [51–60]; however, a direct relation to VCI was not found previously. SOD2, also known as manganese (Mn) dependent Superoxide Dismutase, is located in the mitochondrial matrix and is an enzyme protective against OS [61]. SOD2 was linked to cognitive deficits after mild acute ischemic stroke in early 2022 [43]. Similar to SOD2, SLC30A10, and ATP13A2, are related to Mn [62, 63], and Mn is an important element for the neuronal health [64, 65], which can cross the blood-brain barrier (BBB) [66]. Furthermore, dysfunction of Mn-related proteins (ATP13A2, SLC30A10, SOD2) can increase neuronal damage in stress contexts. Furthermore, the VKORC1 and ATP13A2 proteins in this table were also identified as critical OS-related proteins in the Global Network analysis.

Another protein identified in our study as a VCI-relevant protein, HMGB1, is a nuclear protein released upon inflammasome activation [67]. Previous research has proposed HMGB1 to be a biomarker of cognitive dysfunction [68] and a biomarker of Alzheimer’s disease [69]. Moreover, HMGB1 was proposed to be related to CVD conditions [70–72]. AN, V. et al., [73] suggested that HMGB1 might be a potential target for VCI. Two subunits of Nfk-b (REL and RELA), FLNA and VKORC1 were previously related to VCI [74–76]. LMNA, while not associated with VCI, was previously related to CD and CVD in separate studies [77, 78]. The results shown in **Supplementary Table S6** indicate that 9 proteins that we identified (DOLK, TSC1, ATP1A1, MAPK14, YWHAZ, CREB3, HSPB1, PRDX6, LMNA) have the potential to be novel VCI-relevant proteins as they were not to our knowledge previously linked to VCI in the literature; however, studies demonstrated their link to both CVD and CD. These findings fit with published reports suggesting that dysregulation of Mn produces cognitive impairment [79].

### The Potential Role and Effect of OS

The topological analysis on the Global Network (**Fig. 4**) and the scoring analysis resulted in mostly OS-related proteins in the crosstalk of CVD and CD (**Table 1**). Which led us to focus not only on central proteins but also on OS-related proteins for the crosstalk. The difference between the OS-included and OS-excluded scores of the following 12 proteins with high centralities changed significantly: ATXN1, NEDD4, ATP13A2, VKORC1, FLNA, AKT1, HSPB1, PRDX6, APP, LMNA, PRKN, PSEN1. Of these 12 central OS proteins, five are only OS-related, 1 is CD and OS related, 3 are CVD, CD, and OS related, and 3 are CVD and OS-related proteins. This observation suggests that the central OS proteins are linked more to CVD than CD.

OS plays a crucial role in cell damage during aging [80], and we have found that vascular and neuronal cells are particularly sensitive to OS (**Fig. 6**). Experiments demonstrated that cardiofibroblasts (heart cells) are more resistant to OS. Furthermore, **Supplementary Table S7** shows that, as expected, a large proportion of the OS-related proteins are positively affected when their corresponding OS scores are included in the average calculation. However, our analysis demonstrates that CVD-related proteins are not affected as negatively as CD when OS is included. Interestingly, more CVD-related proteins are affected negatively in total. APP is one of the proteins in that the OS-included score is larger than the OS-excluded score, implying that APP is one of the proteins that OS plays a crucial role in CVD and CD crosstalk.

### Potential Implications of the Identified Pathways

To better understand the roles of 12 OS-significant central proteins in the CVD-CD crosstalk, we investigated which pathways these 12 proteins, their CVD interactors, CD interactors, and CVD & CD interactors were involved in (**Fig. 5, Supplementary Table S5).** HIF1 signaling is one of the enriched pathways. This pathway is related to oxygen homeostasis and the Notch signaling pathway [81]. It was previously shown that HIF1 and Notch signaling pathways are related to VCI [82]. A major result from the CVD interactor enrichment analysis was that two diabetes-related pathways were significantly enriched. Furthermore, for the 3rd enrichment, NRG1 protein on the ERBB4 pathway was previously related to the vascular cognitive disorders [83] (**Supplementary Table S5)**.

The AGE/RAGE signaling pathway was another enriched pathway. AGE/RAGE signaling increases OS leading to diabetes mediated vascular calcification [84]. Previous research linked the AGE/RAGE pathway to CD [85], CVD [86], and VCI [87]. The link between the Notch pathway to CD [88], CVD [89], and VCI [90] are also recognized. We have demonstrated that glycative stress damages neurons (**Supplementary Fig. S3**); however, no research directly linked the AGE/RAGE pathway to CVD and CD crosstalk and its relation to the Notch pathway. Crosstalk between these pathways could play an important role in VCI. A study demonstrated that AGE induced Notch1 signaling led to the inhibition of MMP9 activation causing diabetic wound closure [91]. Another study demonstrated that AGEs induce Notch activation in podocytes that lead to epithelial to mesenchymal transition [92]. Furthermore, C, D.T., et al., [93] indicate that BBB can be damaged by endothelial/epithelial to mesenchymal transition. Therefore, we hypothesize that in the crosstalk between CVD and CD, AGE/RAGE and Notch signaling pathways may be leading to endothelial/epithelial to mesenchymal transition which can be causing BBB disruption. This disruption can lead to BBB leakage and deposition of amyloid beta causing neuronal damage (**Fig. 7**), which is interesting because when the BBB integrity is damaged, it has been proposed that amyloid protein is deposited to repair the holes. Additionally, upregulated RAGE expression promotes the transfer of amyloid beta chains from blood to the brain [33, 94].

**Figure 7.**
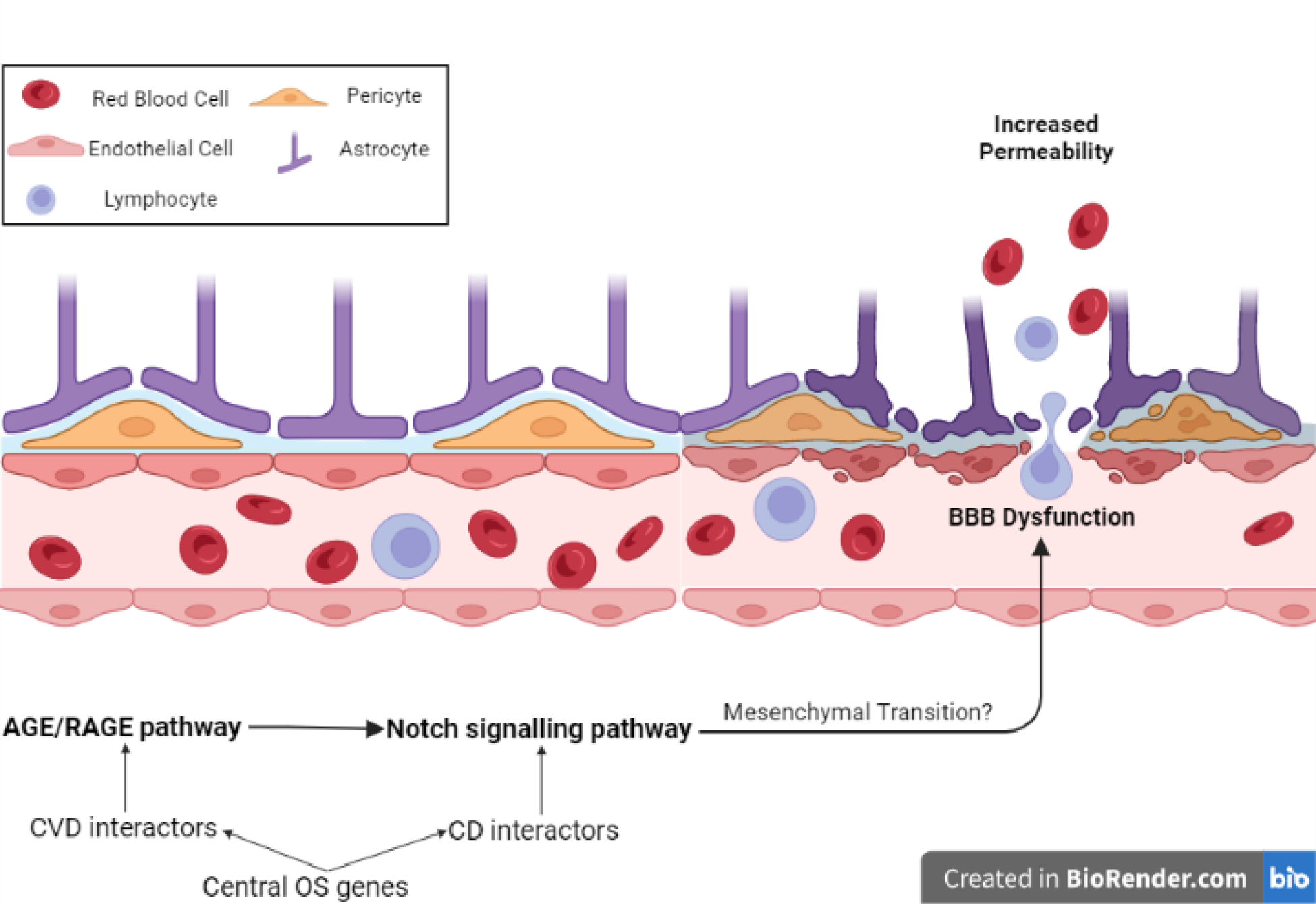
Hypothesized crosstalk of AGE/RAGE and Notch pathways in VCI. Crosstalk between the AGE/RAGE and Notch signaling pathways has been observed in the literature; however, this crosstalk had not previously been linked to VCI. Further, research has shown that the Notch signaling pathway may cause mesenchymal transition; nevertheless, its relation to BBB with this path was not shown. Therefore, we hypothesize that the crosstalk between AGE/RAGE pathway and Notch signaling pathway may be causing mesenchymal transition leading to BBB damage and leakage, causing increased permeability. By the loss of selectivity in BBB, OS will increase. The increased OS levels can elevate the amyloid beta production in the brain. This mechanism as a whole may be causing vascular originated cognitive degeneration. Created with BioRender.com.

## Conclusions and Future Directions

A crosstalk between the AGE/RAGE and Notch signaling pathways might be crucial for VCI. A wide range of previously unrecognized proteins could play a key role in the onset and progression of VCI due to their implication in diseases affecting the cardiovascular system. Heart, brain, and vessel cells appear to have different sensitivities to OS, and glycative stress damages neurons. Our work provides a basis for further work seeking to identify possible therapeutic targets and VCI biomarkers.

## Acknowledgements.

This project has been funded in part by TUBITAK Research Grant No: 220N252 and European Research Area Net (ERANET) ERA-CVD_JTC2020-015 (AG); the Spanish Institute of Health Carlos III by project reference AC20/00009 and AC20/00001 -FEDER/UE and European Research Area Net (ERANET) ERA-CVD_JTC2020-015 (JGO); the “María de Maeztu Programme” for Units of Excellence in Research and Development (R&D, award CEX2018-000792-M); the Spanish Ministry of Science and Innovation and Agencia Estatal de Investigación plus European Regional Development Fund (FEDER Funds) through grants PID2020-117691RB- I00/AEI/10.13039/501100011033 (FJM), PID2020-113203RB-I00 (BO) and PGC2018-101251- B-I00 (JGO).

## Supporting Information

Additional experimental details. Supplementary Figure S1: Count of seed proteins in each subphenotype; Supplementary Figure S2: OS-included and OS-excluded score distribution for the proteins in the Global network; Supplementary Figure S3. Methylglyoxal (MG) neurotoxic effect in mouse cortical primary cultures; Supplementary Figure S4: The node-degree distribution of the global network. Supplementary Table S1 and Supplementary Table S2: Results of overlaps between pairs of CVD and CD subphenotypes, CVD and OS and between CD and OS subphenotypes; Supplementary Table S3: Global Network Analysis Results; Supplementary Table S4 and Supplementary Table S5: Pathway Enrichment Results; Supplementary Table S6: Potentially relevant proteins and their relation to VCI, CVD and CD based on a literature search; Supplementary Table S7: Number of proteins per category (in percentage), the effect of OS to their average GUILD Scores.

## Author Contributions

M.E.Z., S.S., F.J.M.L., B.O., J.G.-O., E.B. R.V., A. J. O. S., A. P. P., A.G., and O.K. designed and conceptualized the project. M.E.Z. and S.S. analyzed data, prepared tables and figures. P.P.P, A.G.E. and M.T. performed cell experiments. M.E.Z., S.S., A.G., O.K., F.J.M.L., B.O., E.B. S.N., A. J. O. S., and A. P. P. wrote and edited the manuscript. E.B., A. J. O. S., A. P. P., provided the Disease-Phenotype taxonomy. All of the authors reviewed and approved the final manuscript.

## Data Availability

The datasets generated during and/or analyzed during the current study are available from the corresponding author on reasonable request.

## References.

1. M, D. and L. D, Vascular Cognitive Impairment. Circulation research, 2017. 120(3).

2. T, I. and S. R, Protein networks in disease. Genome research, 2008. 18(4).

3. AL, B. and O. ZN, Network biology: understanding the cell’s functional organization. Nature reviews. Genetics, 2004. 5(2).

4. X, H. and Z. J, Why do hubs tend to be essential in protein networks? PLoS genetics, 2006. 2(6).

5. U, K. and E. A, Protein-protein interaction networks: probing disease mechanisms using model systems. Genome medicine, 2013. 5(4).

6. J, X. and L. Y, *Discovering disease-genes by topological features in human protein-protein interaction network.* Bioinformatics (Oxford, England), 2006. 22(22).

7. Zuo, W. and J. Wu, The interaction and pathogenesis between cognitive impairment and common cardiovascular diseases in the elderly. https://doi.org/10.1177/20406223211063020, 2022.

8. EE, M. and J. AL, Impact of Cardiovascular Hemodynamics on Cognitive Aging. Arteriosclerosis, thrombosis, and vascular biology, 2021. 41(4).

9. L, P., et al., The role of inflammasomes in vascular cognitive impairment. Molecular neurodegeneration, 2022. 17(1).

10. CM, S., et al., Vascular Oxidative Stress: Impact and Therapeutic Approaches. Frontiers in physiology, 2018. 9.

11. F, C., Chemical Basis of Reactive Oxygen Species Reactivity and Involvement in Neurodegenerative Diseases. International journal of molecular sciences, 2019. 20(10).

12. H, S., B. C, and J. DP, Oxidative Stress. Annual review of biochemistry, 2017. 86.

13. G, I.-R., et al., Amyloid-β peptide fibrils induce nitro-oxidative stress in neuronal cells. Journal of Alzheimer’s disease : JAD, 2010. 22(2).

14. Princeton University “About WordNet.” WordNet. Princeton University. 2010.

15. J, A.-P., et al., GUILDify v2.0: A Tool to Identify Molecular Networks Underlying Human Diseases, Their Comorbidities and Their Druggable Targets. Journal of molecular biology, 2019. 431(13).

16. J, G.-G., et al., Biana: a software framework for compiling biological interactions and analyzing networks. BMC bioinformatics, 2010. 11.

17. J, P., et al., The DisGeNET knowledge platform for disease genomics: 2019 update. Nucleic acids research, 2020. 48(D1).

18. Consortium, U., UniProt: the universal protein knowledgebase in 2021. Nucleic acids research, 2021. 49(D1).

19. J, A., et al., McKusick’s Online Mendelian Inheritance in Man (OMIM). Nucleic acids research, 2009. 37(Database issue).

20. A, C.-a., et al., MINT: the Molecular INTeraction database. Nucleic acids research, 2007. 35(Database issue).

21. I, X., et al., DIP: the database of interacting proteins. Nucleic acids research, 2000. 28(1).

22. S, O., et al., The MIntAct project--IntAct as a common curation platform for 11 molecular interaction databases. Nucleic acids research, 2014. 42(Database issue).

23. Hagberg, A.A., D.A. Schult, and P. Swart. Exploring Network Structure, Dynamics, and Function using NetworkX. 2008.

24. P, S., et al., Cytoscape: a software environment for integrated models of biomolecular interaction networks. Genome research, 2003. 13(11).

25. M, H. and N. S, Implications of Inflammation and Fibrosis in Atrial Fibrillation Pathophysiology. Cardiac electrophysiology clinics, 2021. 13(1).

26. L, Y., X. J, and N. S, Molecular determinants of cardiac fibroblast electrical function and therapeutic implications for atrial fibrillation. Cardiovascular research, 2011. 89(4).

27. M, T., et al., Methylglyoxal reduces mitochondrial potential and activates Bax and caspase-3 in neurons: Implications for Alzheimer’s disease. Neuroscience letters, 2014. 580.

28. E, R.-F., et al., Posttranslational nitro-glycative modifications of albumin in Alzheimer’s disease: implications in cytotoxicity and amyloid-β peptide aggregation. Journal of Alzheimer’s disease : JAD, 2014. 40(3).

29. Y, Z., et al., *Differential Targeting of SLC30A10/ZnT10 Heterodimers to Endolysosomal Compartments Modulates EGF-Induced MEK/ERK1/2 Activity.* Traffic (Copenhagen, Denmark), 2016. 17(3).

30. Hosford, P.S., et al., CO2 signaling mediates neurovascular coupling in the cerebral cortex. Nature Communications, 2022. 13(1): p. 1–11.

31. H, C., et al., Blood Neuroexosomal Mitochondrial Proteins Predict Alzheimer Disease in Diabetes. Diabetes, 2022. 71(6).

32. E, T., et al., Increased intrathecal levels of the angiogenic factors VEGF and TGF-beta in Alzheimer’s disease and vascular dementia. Neurobiology of aging, 2002. 23(2).

33. B, H., F. C, and C. J, Blood-Brain Barrier Breakdown: An Emerging Biomarker of Cognitive Impairment in Normal Aging and Dementia. Frontiers in neuroscience, 2021. 15.

34. RB, H., et al., Cardiovascular manifestations of tuberous sclerosis complex and summary of the revised diagnostic criteria and surveillance and management recommendations from the International Tuberous Sclerosis Consensus Group. Journal of the American Heart Association, 2014. 3(6).

35. PF, K., et al., The Influence of Na(+), K(+)-ATPase on Glutamate Signaling in Neurodegenerative Diseases and Senescence. Frontiers in physiology, 2016. 7.

36. M, P., et al., Tsc1 (hamartin) confers neuroprotection against ischemia by inducing autophagy. Nature medicine, 2013. 19(3).

37. M, H., et al., A systematic analysis of genetic dilated cardiomyopathy reveals numerous ubiquitously expressed and muscle-specific genes. European journal of heart failure, 2015. 17(5).

38. B, S., et al., *Soluble adenylyl cyclase in vascular endothelium: gene expression control of epithelial sodium channel-α, Na+/K+-ATPase-α/β, and mineralocorticoid receptor.* Hypertension (Dallas, Tex. : 1979), 2014. 63(4).

39. A, H., et al., Dolichol kinase deficiency (DOLK-CDG) with a purely neurological presentation caused by a novel mutation. Molecular genetics and metabolism, 2013. 110(3).

40. Z, S., et al., PARK2-dependent mitophagy induced by acidic postconditioning protects against focal cerebral ischemia and extends the reperfusion window. Autophagy, 2017. 13(3).

41. YJ, K., et al., Protective effects of APOE e2 against disease progression in subcortical vascular mild cognitive impairment patients: A three-year longitudinal study. Scientific reports, 2017. 7(1).

42. W, L., et al., Dl-3-n-Butylphthalide Alleviates Hippocampal Neuron Damage in Chronic Cerebral Hypoperfusion via Regulation of the CNTF/CNTFRα/JAK2/STAT3 Signaling Pathways. Frontiers in aging neuroscience, 2021. 12.

43. MS, Z., et al., Low Serum Superoxide Dismutase Is Associated With a High Risk of Cognitive Impairment After Mild Acute Ischemic Stroke. Frontiers in aging neuroscience, 2022. 14.

44. MC, R., T. C, and I.-A. ML, Emerging molecular mechanisms of vascular dementia. Current opinion in hematology, 2019. 26(3).

45. L, B. and C. A, Vascular cognitive disorder. A biological and clinical overview. Neurochemical research, 2010. 35(12).

46. J, H., et al., Quercetin targets VCAM1 to prevent diabetic cerebrovascular endothelial cell injury. Frontiers in aging neuroscience, 2022. 14.

47. J, H., et al., Vascular Cognitive Impairment: Information from Animal Models on the Pathogenic Mechanisms of Cognitive Deficits. International journal of molecular sciences, 2019. 20(10).

48. I, P., et al., PSEN1 Compound Heterozygous Mutations Associated with Cerebral Amyloid Angiopathy and Cognitive Decline Phenotype. International journal of molecular sciences, 2021. 22(8).

49. G, L., et al., ALDOA protects cardiomyocytes against H/R-induced apoptosis and oxidative stress by regulating the VEGF/Notch 1/Jagged 1 pathway. Molecular and cellular biochemistry, 2021. 476(2).

50. D, M., et al., The alterations of Ca2+/calmodulin/CaMKII/CaV1.2 signaling in experimental models of Alzheimer’s disease and vascular dementia. Neuroscience letters, 2013. 538.

51. X, L., et al., Bmal1 Regulates the Redox Rhythm of HSPB1, and Homooxidized HSPB1 Attenuates the Oxidative Stress Injury of Cardiomyocytes. Oxidative medicine and cellular longevity, 2021. 2021.

52. UA, G. and A. JJ, P38α MAPK Signaling-A Robust Therapeutic Target for Rab5-Mediated Neurodegenerative Disease. International journal of molecular sciences, 2020. 21(15).

53. SJ, J., P. JG, and O. GT, *Peroxiredoxins as Potential Targets for Cardiovascular Disease.* Antioxidants (Basel, Switzerland), 2021. 10(8).

54. Q, G., et al., Downregulation of 14-3-3 Proteins in Alzheimer’s Disease. Molecular neurobiology, 2020. 57(1).

55. NA, T. and B. NM, Cardiac Fibroblast p38 MAPK: A Critical Regulator of Myocardial Remodeling. Journal of cardiovascular development and disease, 2019. 6(3).

56. M, H., et al., Neuropathy-causing mutations in HSPB1 impair autophagy by disturbing the formation of SQSTM1/p62 bodies. Autophagy, 2019. 15(6).

57. L, S., D.G. P, and A. C, CREB3 Transcription Factors: ER-Golgi Stress Transducers as Hubs for Cellular Homeostasis. Frontiers in cell and developmental biology, 2019. 7.

58. JH, Q., et al., Proteomic Landscape and Deduced Functions of the Cardiac 14-3-3 Protein Interactome. Cells, 2022. 11(21).

59. HA, K. and M. CE, The Role of Mammalian Creb3-Like Transcription Factors in Response to Nutrients. Frontiers in genetics, 2019. 10.

60. A, K., et al., Overexpression of wild-type human amyloid precursor protein alters GABAergic transmission. Scientific reports, 2021. 11(1).

61. FR, P., et al., Mitochondrial Superoxide Dismutase: What the Established, the Intriguing, and the Novel Reveal About a Key Cellular Redox Switch. Antioxidants & redox signaling, 2020. 32(10).

62. S, A. and T. K, Genetic Disorders of Manganese Metabolism. Current neurology and neuroscience reports, 2019. 19(6).

63. M, L., et al., Zinc transporter 10 (ZnT10)-dependent extrusion of cellular Mn2+ is driven by an active Ca2+-coupled exchange. The Journal of biological chemistry, 2019. 294(15).

64. KJ, H., et al., Manganese Is Essential for Neuronal Health. Annual review of nutrition, 2015. 35.

65. DS, H., et al., Manganese-Induced Neurotoxicity: New Insights Into the Triad of Protein Misfolding, Mitochondrial Impairment, and Neuroinflammation. Frontiers in neuroscience, 2019. 13.

66. J, B., et al., Impact of manganese on and transfer across blood-brain and blood-cerebrospinal fluid barrier in vitro. The Journal of biological chemistry, 2012. 287(21).

67. R, C., K. R, and T. D, The mechanism of HMGB1 secretion and release. Experimental & molecular medicine, 2022. 54(2).

68. YN, P., et al., HMGB1: A Common Biomarker and Potential Target for TBI, Neuroinflammation, Epilepsy, and Cognitive Dysfunction. Frontiers in neuroscience, 2018. 12.

69. BW, F., et al., HMGB1 and thrombin mediate the blood-brain barrier dysfunction acting as biomarkers of neuroinflammation and progression to neurodegeneration in Alzheimer’s disease. Journal of neuroinflammation, 2016. 13(1).

70. W, L., S. AE, and W. H, Role of HMGB1 in cardiovascular diseases. Current opinion in pharmacology, 2006. 6(2).

71. J, C., et al., The Role of HMGB1 in Cardiovascular Biology: Danger Signals. Antioxidants & redox signaling, 2015. 23(17).

72. A, R., et al., The Janus face of HMGB1 in heart disease: a necessary update. Cellular and molecular life sciences : CMLS, 2019. 76(2).

73. AN, V., et al., Role of HMGB1 in an Animal Model of Vascular Cognitive Impairment Induced by Chronic Cerebral Hypoperfusion. International journal of molecular sciences, 2020. 21(6).

74. R, S., et al., Astroglial NF-kB contributes to white matter damage and cognitive impairment in a mouse model of vascular dementia. Acta neuropathologica communications, 2016. 4(1).

75. J, M., et al., Variation in VKORC1 Is Associated with Vascular Dementia. Journal of Alzheimer’s disease : JAD, 2021. 80(3).

76. C, B., et al., Cardiovascular and connective tissue disorder features in FLNA-related PVNH patients: progress towards a refined delineation of this syndrome. Orphanet journal of rare diseases, 2021. 16(1).

77. S, C., M. I, and D.P. E, The Broad Spectrum of LMNA Cardiac Diseases: From Molecular Mechanisms to Clinical Phenotype. Frontiers in physiology, 2020. 11.

78. B, F., B. FH, and F. MB, Lamin Dysfunction Mediates Neurodegeneration in Tauopathies. Current biology : CB, 2016. 26(1).

79. RC, B., et al., Brain manganese and the balance between essential roles and neurotoxicity. The Journal of biological chemistry, 2020. 295(19).

80. I, L., et al., Oxidative stress, aging, and diseases. Clinical interventions in aging, 2018. 13.

81. X, Z., et al., Interaction with factor inhibiting HIF-1 defines an additional mode of cross-coupling between the Notch and hypoxia signaling pathways. Proceedings of the National Academy of Sciences of the United States of America, 2008. 105(9).

82. YL, C., et al., Evidence that collaboration between HIF-1α and Notch-1 promotes neuronal cell death in ischemic stroke. Neurobiology of disease, 2014. 62.

83. Y, H., et al., Neuregulin1 attenuates cognitive deficits and hippocampal CA1 neuronal apoptosis partly via ErbB4 receptor in a rat model of chronic cerebral hypoperfusion. Behavioural brain research, 2019. 365.

84. AM, K. and S. JA, RAGE Differentially Altered in vitro Responses in Vascular Smooth Muscle Cells and Adventitial Fibroblasts in Diabetes-Induced Vascular Calcification. Frontiers in physiology, 2021. 12.

85. J, C., et al., Assessment of Advanced Glycation End Products and Receptors and the Risk of Dementia. JAMA network open, 2021. 4(1).

86. F, S., et al., Effects of RAGE Deletion on the Cardiac Transcriptome during Aging. International journal of molecular sciences, 2022. 23(19).

87. L, S., W. J, and E. MM, Immunohistochemical study of N-epsilon-carboxymethyl lysine (CML) in human brain: relation to vascular dementia. BMC neurology, 2007. 7.

88. SJ, C., et al., Altered expression of Notch1 in Alzheimer’s disease. PloS one, 2019. 14(11).

89. HE, M., et al., Notch3 signalling and vascular remodelling in pulmonary arterial hypertension. Clinical science (London, England : 1979), 2019. 133(24).

90. A, K. and N. DA, Role of Notch signaling in neurovascular aging and Alzheimer’s disease. Seminars in cell & developmental biology, 2021. 116.

91. P, Z., et al., AGEs-induced MMP-9 activation mediated by Notch1 signaling is involved in impaired wound healing in diabetic rats. Diabetes research and clinical practice, 2022. 186.

92. R, N., et al., Activation of Notch1 signaling in podocytes by glucose-derived AGEs contributes to proteinuria. BMJ open diabetes research & care, 2020. 8(1).

93. C, D.T., et al., Molecular alterations of the blood-brain barrier under inflammatory conditions: The role of endothelial to mesenchymal transition. Biochimica et biophysica acta, 2016. 1862(3).

94. R, D., et al., RAGE mediates amyloid-beta peptide transport across the blood-brain barrier and accumulation in brain. Nature medicine, 2003. 9(7).

