## Supplementary Information for "Revealing Shared Proteins and Pathways in Cardiovascular and Cognitive Diseases Using Protein Interaction Network Analysis"

#### Supplementary Figures

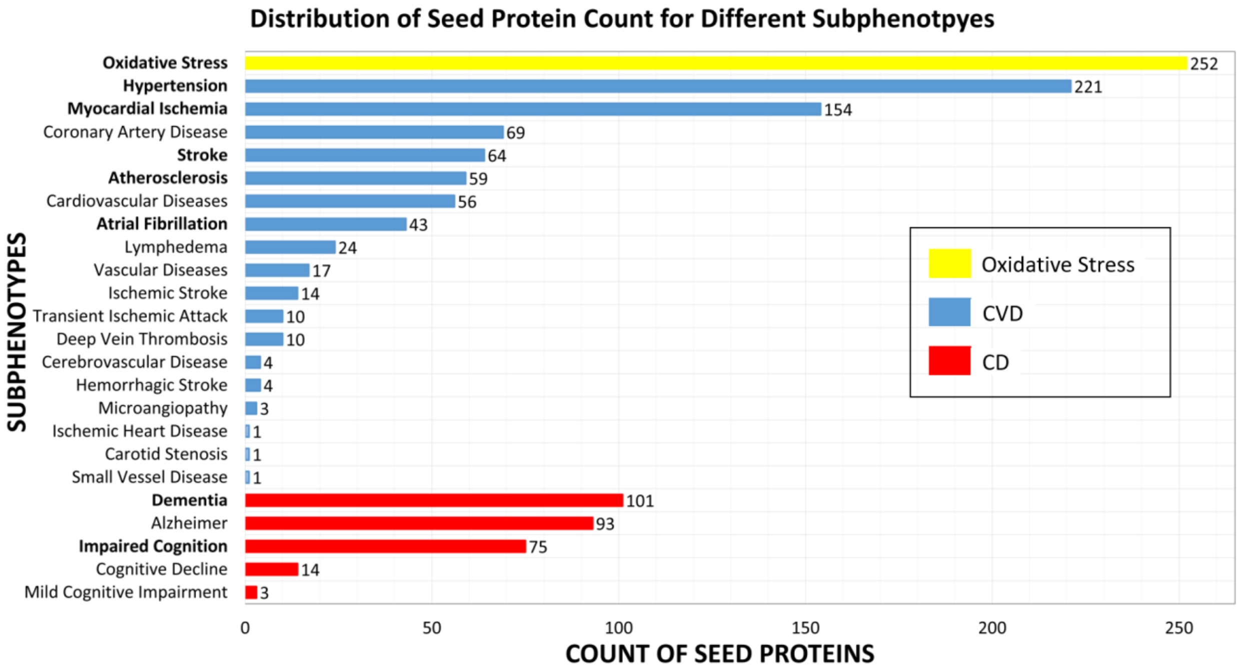

**Supplementary Figure S1**. Count of seed proteins in each subphenotype. The bars are colored according to the subphenotype category: Yellow for OS, blue for Cardiovascular Diseases (CVD) and red for Cognitive Diseases (CD). Selected subphenotypes to represent CVD or CD are highlighted in bold.

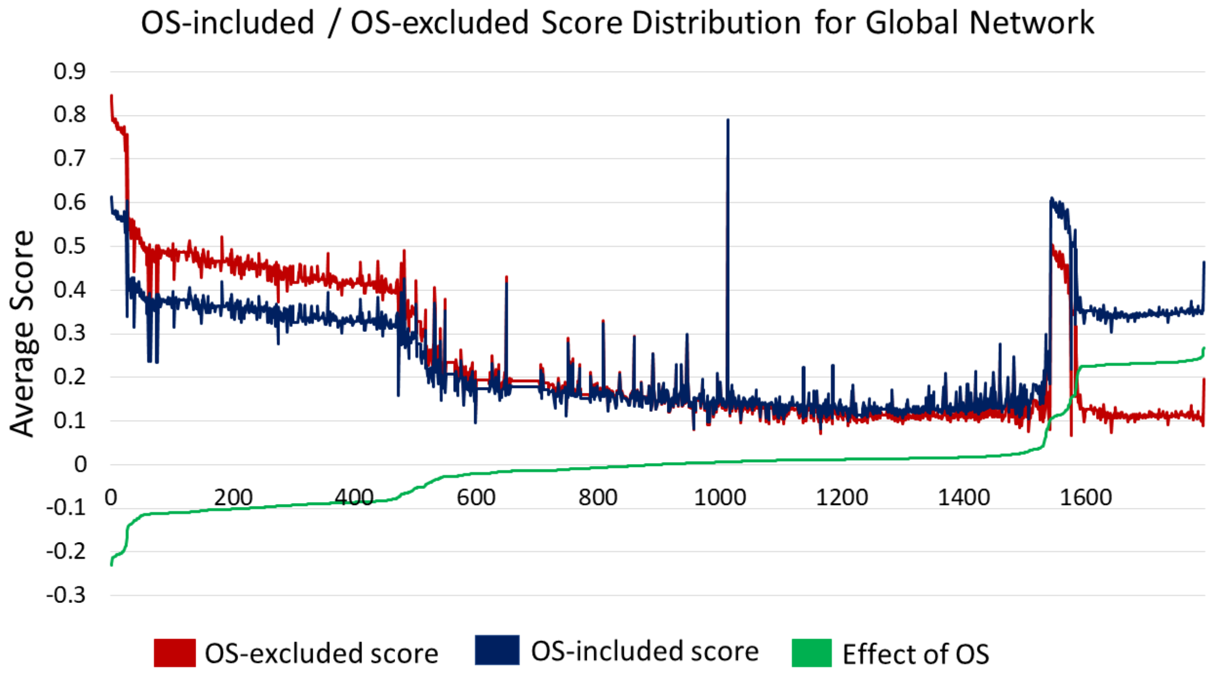

**Supplementary Figure S2.** OS-included and OS-excluded score distribution for the proteins in the Global network. The y-axis is the average score, and the x axis represents the proteins, scored in terms of their average score (OS-included/OS-excluded and their difference given as effect of OS). Results indicated that out of the 1796 proteins in the global network, 893 of them are negatively affected by OS (decreased OS-excluded score), and 903 of them are positively affected (increased OS-excluded score).

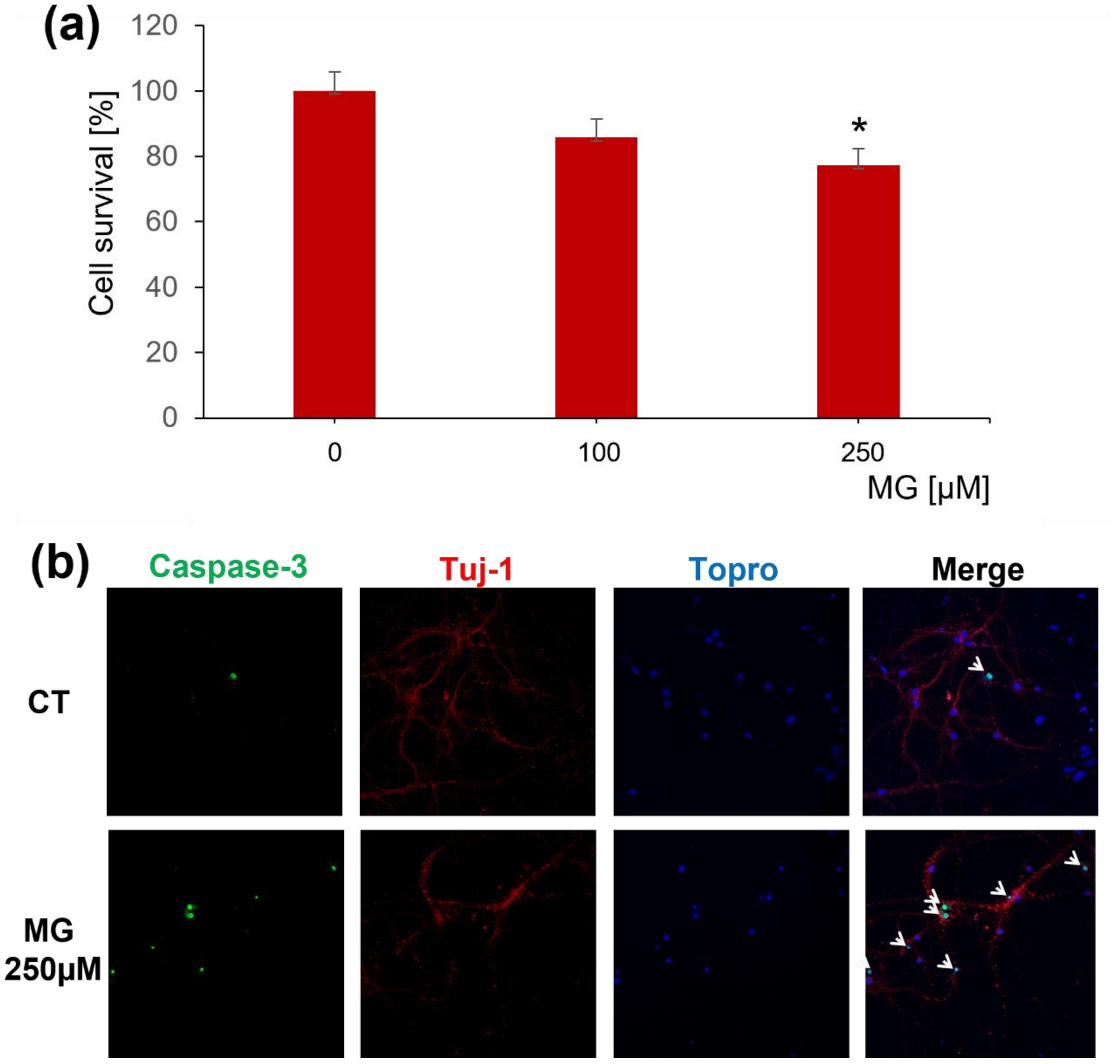

**Supplementary Figure S3**. Methylglyoxal (MG) neurotoxic effect in mouse cortical primary cultures. (**a**) Cells challenged with increasing concentrations of MG and assayed by MTT reduction. Data are mean ± SEM of 6 independent experiments performed in triplicate. * $p<0.05$ *vs* the respective controls by ANOVA plus Tukey-Kramer Multiple Comparisons Test. (**b**) Immunofluorescence study of cortical primary cultures treated with 250 μM MG for 24 h looking for caspase-3 activation (green) and microtubule integrity (red). Nuclei are stained with Topro (blue). White arrows show the localization of Caspase-3 in cell soma.

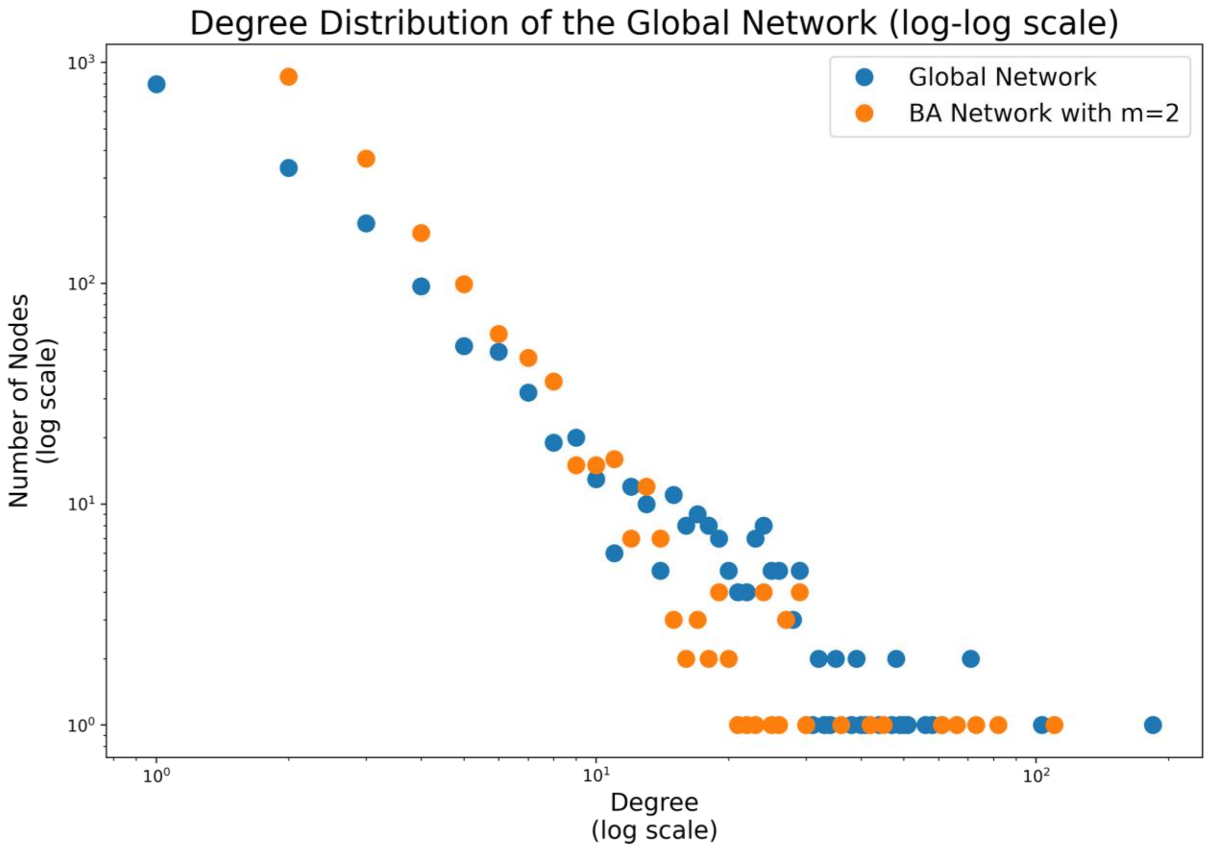

**Supplementary Figure S4**. The node-degree distribution of the global network on a log-log scale. The BA (Barabasi Albert) network represents the degree distribution of a random scale-free BA network with the same number of nodes as the global network. The global network has a similar distribution with the BA network. “m=2” represents the number of preferential attachments a newly included node makes.

### Supplementary Tables

**Supplementary Table S1**. Results of overlaps between pairs of CVD and CD subphenotypes[1].

| Subphenotype 1 | Subphenotype 2 | Protein | Score |
|---|---|---|---|
| Atherosclerosis (CVD) | Dementia (CD) | SERPINI1 (Neuroserpin)<br>CLN6 (Ceroid-lipofuscinosis neuronal protein 6)<br>**VEGFA** (Vascular endothelial growth factor A) | 0.76<br>0.51<br>0.39 |
| Atrial Fibrillation (CVD) | Dementia (CD) | **CLN8** (Protein CLN8)<br>**ATP13A2** (Polyamine-transporting ATPase 13A2)<br>**LMNA** (Prelamin-A/C) | 0.48<br>0.48<br>0.44 |
| Hypertension (CVD) | Dementia (CD) | **VEGFA** (Vascular endothelial growth factor A)<br>**SMAD4** (Mothers against decapentaplegic homolog 4)<br>NRP2 (Neuropilin-2) | 0.46<br>0.42<br>0.21 |
| Myocardial Ischemia (CVD) | Dementia (CD) | **VEGFA**(Vascular endothelial growth factor A)<br>**HMGB1** (High mobility group protein B1)<br>NRP2 (Neuropilin-2) | 0.51<br>0.45<br>0.30 |
| Stroke (CVD) | Dementia (CD) | **ATXN1** (Ataxin-1)<br>**RELA** (Transcription factor p65)<br>**REL** (Proto-oncogene c-Rel) | 0.45<br>0.40<br>0.40 |
| Atherosclerosis (CVD) | Impaired Cognition (CD) | GJB2 (Gap junction beta-2 protein)<br>**DOLK** (Dolichol kinase)<br>**FLNA** (Filamin-A) | 0.47<br>0.45<br>0.45 |
| Atrial Fibrillation (CVD) | Impaired Cognition (CD) | DOLK (Dolichol kinase)<br>GJB2 (Gap junction beta-2 protein)<br>**FLNA** (Filamin-A) | 0.52<br>0.52<br>0.46 |
| Hypertension (CVD) | Impaired Cognition (CD) | **TSC1** (Hamartin)<br>**KRT8** (Keratin, type II cytoskeletal 8)<br>**SH2B1** (SH2B adapter protein 1) | 0.44<br>0.43<br>0.12 |
| Myocardial Ischemia (CVD) | Impaired Cognition (CD) | COL5A1 (Collagen alpha-1(V) chain)<br>**ATP1A1** (Sodium/potassium-transporting ATPase subunit alpha-1)<br>**KMT2C** (Histone-lysine N-methyltransferase 2C) | 0.47<br>0.43<br>0.12 |
| Stroke (CVD) | Impaired Cognition (CD) | **FLNA** (Filamin-A)<br>**SDHB** (Succinate dehydrogenase [ubiquinone] iron-sulfur subunit, mitochondrial)<br>**ATXN1** (Ataxin-1) | 0.44<br>0.44<br>0.43 |

[1] Top 3 overlap scores of non-seed and linker genes are shown, highlighting in bold the linker proteins with the corresponding protein names in parentheses. Overlap scores are shown in the last column.

**Supplementary Table S2.** Results of overlaps between pairs of CVD and OS and between CD and OS subphenotypes[2].

| Subphenotype | OS | Protein | Overlap Score |
|---|---|---|---|
| Atherosclerosis (CVD) | Oxidative Stress | **MMP2** (72 kDa type IV collagenase)<br>**UBQLN1** (Ubiquilin-1)<br>**TP53** (Cellular tumor antigen p53) | 0.45<br>0.44<br>0.43 |
| Atrial Fibrillation (CVD) | Oxidative Stress | MSRB1 (Methionine-R-sulfoxide reductase B1)<br>SLC30A10 (Zinc transporter 10)<br>**VKORC1** (Vitamin K epoxide reductase complex subunit 1) | 0.75<br>0.69<br>0.50 |
| Hypertension (CVD) | Oxidative stress | **GTF2I** (General transcription factor II-I)<br>**MYH9** (Myosin-9)<br>DESI1 (Desumoylating isopeptidase 1) | 0.45<br>0.44<br>0.29 |
| Myocardial Ischemia (CVD) | Oxidative stress | MMP3 (Stromelysin-1)<br>**MEOX2** (Homeobox protein MOX-2)<br>**TKT** (Transketolase) | 0.52<br>0.48<br>0.15 |
| Stroke (CVD) | Oxidative stress | **REL** (Proto-oncogene c-Rel)<br>**PPP2CA** (Serine/threonine-protein phosphatase 2A catalytic subunit alpha isoform)<br>**CSNK2A1** (Casein kinase II subunit alpha) | 0.43<br>0.09<br>0.09 |
| Dementia (CD) | Oxidative stress | CYGB (Cytoglobin)<br>PDK2 ([Pyruvate dehydrogenase (acetyl-transferring)] kinase isozyme 2, mitochondrial)<br>**ATP13A2** (Polyamine-transporting ATPase 13A2) | 0.61<br>0.53<br>0.53 |
| Impaired Cognition (CD) | Oxidative stress | **PEX19** (Peroxisomal biogenesis factor 19)<br>GSKIP (GSK3B-interacting protein)<br>ABCD1 (ATP-binding cassette sub-family D member 1) | 0.52<br>0.52<br>0.50 |

[2] Top 3 overlap scores of non-seed and linker genes are shown, highlighting in bold the linker proteins with the corresponding protein names in parentheses. Overlap scores are shown in the last column.

**Supplementary Table S3**. Global Network Analysis Results.

| Property | Results for Global Network |
|---|---|
| Diameter | 13 |
| Average Clustering Coefficient | 0.11 |
| Transitivity | 0.24 |
| Number of Central Genes | 155 |
| Average Shortest Path Length | 4.97 |
| Betweenness Centrality (Top5) | LMNA 0.31<br>ATXN1 0.12<br>REL 0.1<br>FLNA 0.086<br>YWHAZ 0.08 |
| Degree Centrality (Top5) | LMNA 0.11<br>ATXN1 0.06<br>REL 0.041<br>CREB3 0.041<br>HSPB1 0.035 |
| Change in Average Clustering Coefficient when Network Perturbed (Top5) | LMNA 0.0086<br>CREB3 0.0081<br>ATXN1 -0.0023<br>ATXN1L 0.0022<br>MEOX2 -0.0019 |

**Supplementary Table S4.** Pathway enrichment for first set of genes and their interactors[3].

| Category | KEGG | REAC |
|---|---|---|
| **Enrichment 1** 16 CVD, CD, CVD and CD genes + interacting CVD genes (174) | Cocaine addiction | Neurexins and neuroligins |
| | Amphetamine addiction | Neuronal System |
| | Pathways in cancer | Protein-protein interactions at synapses |
| | Chagas disease | Synaptic adhesion-like molecules |
| | AGE-RAGE signaling pathway in diabetic complications | Assembly and cell surface presentation of NMDA receptors |
| **Enrichment 2** 16 CVD, CD, CVD and CD genes + interacting CD genes (31) | Pathways of neurodegeneration - multiple diseases | |
| | Hippo signaling pathway | |
| **Enrichment 3** 16 CVD, CD, CVD and CD genes + interacting CVD and CD genes (10) | Focal adhesion | Signal Transduction |
| | Hippo signaling pathway | Signaling by Receptor Tyrosine Kinases |
| | AGE-RAGE signaling pathway in diabetic complications | Signaling by TGFB family members |
| | PI3K-Akt signaling pathway | - |
| **Enrichment 4** 16 CVD, CD, CVD and CD genes + interacting CVD genes (174) + interacting CD genes (31) + interacting CVD and CD genes (10) | Cocaine addiction | Neurexins and neuroligins |
| | Amphetamine addiction | Protein-protein interactions at synapses |
| | Pathways in cancer | Diseases of signal transduction by growth factor receptors and second messengers |
| | Chagas disease | Synaptic adhesion-like molecules |
| | Fluid shear stress and atherosclerosis | Signal Transduction |

[3] CVD interactors, CD interactors and CVD and CD interactors of ALDOA, RELA, SMAD1, FLOT1, DLG4, SYNE4, YWHAZ, FLNB, SMAD4, APOE, VCAM1, PPP1CA, CAV1, FN1, EWSR1, TCF4.

**Supplementary Table S5**. Pathway enrichment for Central OS genes and their interactors.

| Category | KEGG | REAC |
|---|---|---|
| **CVD Interactors**<br>12 OS proteins<br>+<br>interacting CVD proteins (199)<br>+<br>interacting CVD & OS proteins (31) | AGE-RAGE signaling pathway in diabetic complications | Plasma lipoprotein assembly |
| | HIF-1 signaling pathway | Transport of small molecules |
| | Diabetic cardiomyopathy | Platelet degranulation |
| | Sphingolipid signaling pathway | Response to elevated platelet cytosolic Ca2+ |
| | Central carbon metabolism in cancer | Platelet activation, signaling and aggregation |
| **CD Interactors**<br>12 OS proteins<br>+<br>interacting CD proteins (72)<br>+<br>interacting<br>CD & OS proteins (68) | Pathways of neurodeproteinration - multiple diseases | Amyloid fiber formation |
| | Parkinson disease | Josephin domain DUBs |
| | Notch signaling pathway | NRIF signals cell death from the nucleus |
| | | Diseases of signal transduction by growth factor receptors and second messengers |
| | | Noncanonical activation of NOTCH3 |
| **CVD & CD Interactors**<br>12 OS proteins<br>+<br>interacting CVD & CD proteins (14)<br>+<br>interacting CVD & CD & OS proteins (6) | Alzheimer disease | Platelet activation, signaling and aggregation |
| | Hepatitis C | Signaling by ERBB4 |
| | T cell receptor signaling pathway | Chylomicron clearance |
| | Insulin resistance | Signaling by Receptor Tyrosine Kinases |
| | Neurotrophin signaling pathway | Platelet activation, signaling and aggregation |

**Supplementary Table S6**. Potentially relevant proteins and their relation to VCI, CVD and CD based on a literature search[4].

| Type of Analysis | Protein | CVD | CD | VCI |
|---|---|---|---|---|
| Overlap | VEGFA | Red | Red | Red |
| | DOLK | Red | Red | Green |
| | GJB2 | Red | Red | Red |
| | TSC1 | Red | Red | Green |
| | ATP1A1 | Red | Red | Green |
| | COL5A1 | Red | Red | Red |
| | SDHB | Red | Red | Red |
| | SLC30A10, ZNT10 | Red | Red | Red |
| Global | SOD2 | Red | Red | Red |
| | MAPK14 | Red | Red | Green |
| | JAK2 | Red | Red | Red |
| | YWHAZ | Red | Red | Green |
| | CREB3 | Red | Red | Green |
| | HSPB1 | Red | Red | Green |
| | APOE | Red | Red | Red |
| | ALDOA | Red | Green | Red |
| | VCAM1 | Red | Red | Red |
| | AKT1 | Red | Red | Red |
| | PRDX6 | Red | Red | Green |
| | APP | Red | Red | Red |
| | PSEN1 | Red | Red | Red |
| | NEDD4 | Red | Red | Red |
| | PRKN | Red | Green | Red |
| Overlap and Global | VKORC1 | Red | Green | Red |

[4] As the aim is to identify novel relevant proteins of VCI, if previous research demonstrated a relation to VCI, its individual relation with CVD or CD was not searched. 31 potential proteins were categorized into 3 groups: proteins found crucial only from Overlap analysis, proteins found crucial only from Global analysis, and proteins coming from both analyses. Red boxes represent the proteins with relation to the corresponding column. Green boxes represent the proteins not related to the corresponding column.

| | |
|---|---|
| | ATXN1 |
| | FLNA |
| | HMGB1 |
| | REL |
| | RELA |
| | ATP13A2 |
| | LMNA |

**Supplementary Table S7.** Number of proteins per category (in percentage), the effect of OS to their average GUILD Scores[5].

| | CVD | CD | OS | CVD & CD | CVD & OS | CD & OS | CVD & CD & OS |
|---|---|---|---|---|---|---|---|
| Percentage of proteins affected >= +0.1 | 0 | 0 | 0.44 | 0 | 0.41 | 0.17 | 0.14 |
| Percentage of proteins affected <= -0.1 | 0.08 | 0.37 | 0.09 | 0.36 | 0.19 | 0.06 | 0.14 |
| Percentage of proteins affected <= + 0.1 | 0.37 | 0.38 | 0.40 | 0.21 | 0.23 | 0.25 | 0.29 |
| Percentage of proteins affected 0<= >= - 0.1 | 0.55 | 0.25 | 0.07 | 0.43 | 0.18 | 0.52 | 0.43 |

[5] Columns represent the categories of the proteins. If there is only one category name on a given column, it signifies that a protein only belongs to that category. If a column has more than one category name, the given values represent the number of proteins that are present in both categories. For example, 6% of the proteins (5) shared by CD and OS in which the OS effect is at most -0.1.
